# The behavioral signature of stepwise learning strategy and its neural correlate in the basal forebrain

**DOI:** 10.1101/2022.04.01.486795

**Authors:** Hachi E. Manzur, Ksenia Vlasov, You-Jhe Jhong, Hung-Yen Chen, Shih-Chieh Lin

**Affiliations:** Neural Circuits and Cognition Unit, Laboratory of Behavioral Neuroscience, National Institute on Aging, National Institutes of Health, Baltimore, MD, USA; Institute of Neuroscience, National Yang Ming Chiao Tung University, Taipei, Taiwan; Brain Research Center, National Yang Ming Chiao Tung University, Taipei, Taiwan

## Abstract

**Summary:** Studies of associative learning have commonly focused on how rewarding outcomes are predicted by either sensory stimuli or animals’ actions. However, in many learning scenarios, reward delivery requires the occurrence of both sensory stimuli and animals’ actions in a specific order, in the form of behavioral sequences. How such behavioral sequences are learned is much less understood. Here we provide behavioral and neurophysiological evidence to show that behavioral sequences are learned using a stepwise strategy. In rats learning a new association, learning started from the behavioral event closest to the reward and sequentially incorporated earlier events. This led to the sequential refinement of reward-seeking behaviors, which was characterized by the stepwise elimination of ineffective and non-rewarded behavioral sequences. At the neuronal level, this stepwise learning process was mirrored by the sequential emergence of basal forebrain neuronal responses toward each event, which quantitatively conveyed a reward prediction error signal and promoted reward-seeking behaviors. Together, these behavioral and neural signatures revealed how behavioral sequences were learned in discrete steps and when each learning step took place.

## Introduction

Associative learning is essential for survival and allows animals and humans to predict future reward based on environmental stimuli (*1, 2*) or their own actions (*3*–*5*). Understanding the algorithmic principles of associative learning has been a central question in psychology and neuroscience (*6–13*), and has broad implications in machine learning and artificial intelligence (*14–16*).

While the learning of stimulus-reward and action-reward associations have been historically studied under the separate labels of Pavlovian (*1, 2*) and instrumental (*3, 4*) conditioning, most learning scenarios require the synergistic contribution from both types of learning strategies. For example, when a new reward-predicting stimulus is introduced to the environment, the Pavlovian strategy might not be sufficient because oftentimes the reward would not be delivered unless animals take specific actions. In experimental settings, such actions could be a lever press, or a saccade toward a target, or multiple licks before the reward is delivered. In these scenarios, reward is obtained only when sensory stimuli and animals’ actions occur in a specific order as a behavioral sequence (*5, 17–19*). Compared to the wealth of knowledge about Pavlovian and instrumental conditioning, little is understood about how animals learn behavioral sequences that contain both stimuli and actions.

Converging views from theoretical studies support the idea that reward-predicting behavioral sequences can be efficiently learned using a strategy that we will refer to as stepwise learning: learning starts from the event closest to the reward, while earlier events are learned in later steps. This learning strategy was initially proposed by Skinner (*5*) and more recently elaborated into formal learning models (*18, 19*). Similar learning dynamics are also predicted by reinforcement learning algorithms, in which states that are closer to the final reward are learned first (*14*). The stepwise learning strategy has also been successfully used in various animal training scenarios to incrementally chain single behaviors into long sequences over multiple training steps (*20*).

The goal of the current study is to test whether animals use the stepwise learning strategy to learn reward-predicting behavioral sequences that contain both stimuli and actions. We seek to identify the behavioral and neural signatures of this learning process that can delineate the discrete steps of learning. A major challenge in understanding this type of learning is that behavioral sequences are controlled not only by the experimenter but also by the animal, which is free to take various actions. We reason that, at the beginning of learning, animals’ actions would be less constrained and therefore would generate a large repertoire of behavioral sequences that may or may not lead to the rewarding outcome. As the stepwise learning process unfolds, the repertoire of behavioral sequences should become increasingly selective as well as more frequently rewarded. Therefore, the behavioral signature of the stepwise learning strategy may reside in how the entire repertoire of behavioral sequences become sequentially refined during the learning process. In the current study, we identified such a behavioral signature, which corresponded to the discrete steps in the stepwise learning process.

In order to validate the behavioral signature for the stepwise learning strategy, we focused on a special subset of noncholinergic neurons in the basal forebrain (BF), which are referred to as BF bursting neurons (*21–26*). Previous studies have found that BF bursting neurons convey a reward-prediction error signal (*21, 22, 27, 28*), and show highly robust phasic bursting responses to reward-predicting sensory stimuli irrespective of their sensory modalities (*21–25*). Moreover, such responses only emerge after reward-based associative learning (*21*), and are tightly coupled with behavioral performance and promotes faster decision speeds (*21–23*). These observations suggest that increased BF bursting neuron activities toward a behavioral event reflects that the event has been learned as a reward predictor. By observing the temporal evolution of BF bursting neuron activities throughout the learning process in the current study, we predict that BF responses should mirror the stepwise learning process: BF activity should first emerge toward the last behavioral event closest to the reward, and subsequently develop toward the earlier behavioral events. Such behavioral and neurophysiological findings will provide important insights on how behavioral sequences are learned.

## Results

### A model for learning behavioral sequences using the stepwise learning strategy

To gain intuition about the stepwise learning strategy, we first considered a toy example in which a three-element sequence A-B-C predicted reward (Figure 1A). This sequence can be learned using the stepwise strategy in three discrete steps, starting from the event closest to the reward and sequentially incorporating earlier events (Figure 1B). In each step, behavioral sequences that contained all the learned events in that step would predict reward and therefore preferentially executed, while incompatible sequences would not predict reward and therefore be eliminated from the behavioral repertoire. As a result, the discrete steps of learning would correspond to the stepwise elimination of non-rewarded behavioral sequences that share subsets of behavioral elements (Figure 1B). We hypothesized that this sequential refinement of reward-seeking behaviors might provide a behavioral signature of the stepwise learning strategy.

**Figure 1.**
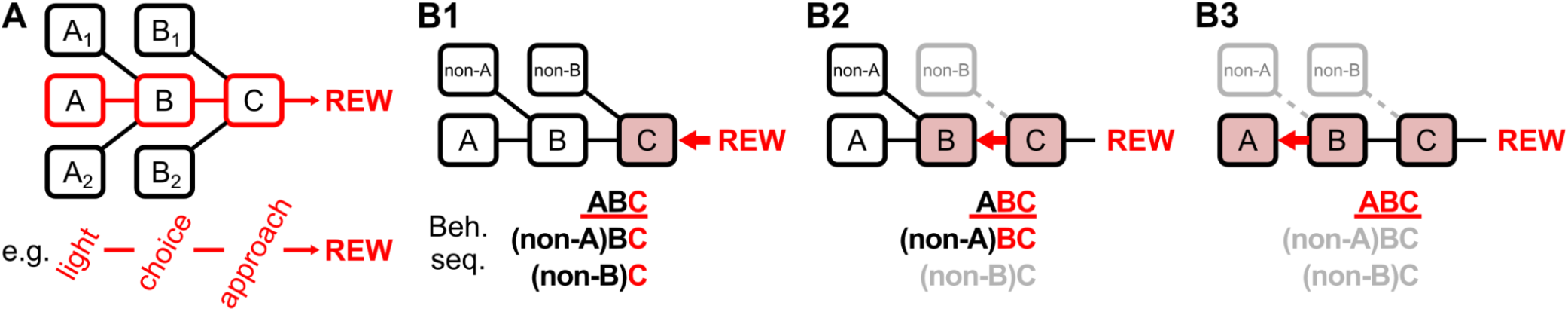
A model for the stepwise learning of behavioral sequences. **A**, Schematic of an example scenario where the reward is predicted by a simple sequence consisting of three behavioral events A-B-C. For example, A could be a light stimulus, B the rightward choice, and C the approach behavior to obtain reward. A1/A2/B1/B2 indicate alternative behavioral elements for A/B that could be combined to generate other behavioral sequences. Among all possible sequences, only the A-B-C sequence is rewarded. **B**, The three distinct steps when learning the A-B-C sequence using the stepwise strategy illustrate how reward-seeking behaviors are sequentially refined. For simplicity, alternative behavioral events are denoted as non-A and non-B. Three possible behavioral sequences are listed. Behavioral events that have been learned as reward-predictors are colored in red, while the rewarded sequence (A-B-C) is underlined in red. **B1**, The first step of learning involves the event closest to the reward, C. Animals would engage in all three behavioral sequences because they all contain this reward-predicting event. **B2**, The second step of learning involves the next-to-last event, B. Behavioral sequences that contain the reward-predicting events B-C are preserved, while the incompatible sequence is eliminated from the behavioral repertoire (gray). **B3**, The third step of learning involves the earliest event, A. Only the A-B-C sequence contains all the reward-predicting events A-B-C and therefore preserved, while another incompatible sequence is eliminated from the behavioral repertoire.

### Sequential refinement of reward-seeking behaviors during new learning

To test whether animals indeed use the stepwise strategy to learn behavioral sequences that contain both sensory stimuli and their own actions, we trained adult Long-Evans rats in an auditory discrimination task. Rats entered the fixation port to initiate each trial, where they encountered three trial types (S^left^; S^right^; catch) with equal probabilities that respectively indicated sucrose water reward in the left or right port, or no reward in the case of catch trials (no stimulus) (Figure 2A). During the initial auditory discrimination phase, S^left^ and S^right^ were two distinct sound stimuli. After reaching asymptotic performance, rats entered the new learning phase (first new learning session denoted as the D_0_ session), in which the S^right^ sound stimulus was switched to a novel light stimulus that minimized sensory generalization from past experience (Figure 2B, Table 1). During the new learning phase, rats maintained stable levels of performance toward the previously-learned S^left^ sound stimulus (Figure 2C) (94.3±4.8% correct, 109±20 trials per session, mean±std). Within the first three sessions of the new learning phase, all rats (N=7) began responding correctly in the new light trials (the first such session denoted as the D_1_ session) and maintained stable levels of >90% correct response rates afterwards (Figure 2C, Table 2).

**Table 1.** Stimulus parameters of the S^left^ and S^right^ stimuli for each animal.

| Animal ID | $S^{\text{left}}$ sound | $S^{\text{right}}$ sound<br>(auditory discrimination) | $S^{\text{right}}$ light<br>(new learning) |
| --- | --- | --- | --- |
| #1 | 12kHz 80dB 2s | 6kHz 80dB 2s | Center Light 0.5s |
| #2 | 100Hz clicker 75dB 1s | white noise 75dB 1s | Center Light 1s |
| #3 | 100Hz clicker 75dB 1s | white noise 75dB 1s | Center Light 1s |
| #4 | 100Hz clicker 75dB 1s | white noise 75dB 1s | Center Light 1s |
| #5 | 100Hz clicker 75dB 1s | white noise 75dB 1s | Center Light 1s |
| #6 | 100Hz clicker 75dB 1s | white noise 75dB 1s | Center Light 1s |
| #7 | 100Hz clicker 75dB 1s | white noise 75dB 1s | Center Light 1s |

**Table 2.** Timing of the three landmark sessions in each animal and the number of recorded BF neurons in each session.

| Sessions (relative to each animal's $D_2$ session) | | | | | | | | |
| --- | --- | --- | --- | --- | --- | --- | --- | --- |
| Animal ID | $D_2 - 3$ | $D_2 - 2$ | $D_2 - 1$ | $D_2$ | $D_2 + 1$ | $D_2 + 2$ | $D_2 + 3$ | $D_2 + 4$ |
| #1 | 4/13 | 1/4 | 23/32 | 25/33 | 34/49 | 28/41 | 31/46 | 29/42 |
| #2 | 26/43 | 29/46 | 26/38 | 25/40 | 25/42 | 13/28 | 15/31 | 18/36 |
| #3 |  | 33/39 | 20/29 | 29/38 | 26/36 | 21/33 | 26/34 | 30/40 |
| #4 |  | 17/27 | 15/21 | 17/23 | 21/30 | 21/25 | 24/26 |  |
| #5 |  |  | 29/35 | 32/39 | 30/38 | 30/39 | 29/36 | 31/35 |
| #6 |  |  | 17/30 | 16/29 | 20/37 | 16/28 | 18/29 | 20/27 |
| #7 |  |  |  | 23/28 | 20/22 | 11/13 | 19/23 |  |

$D_0$ $D_1$ $D_2$ session symbol

**Figure 2.**
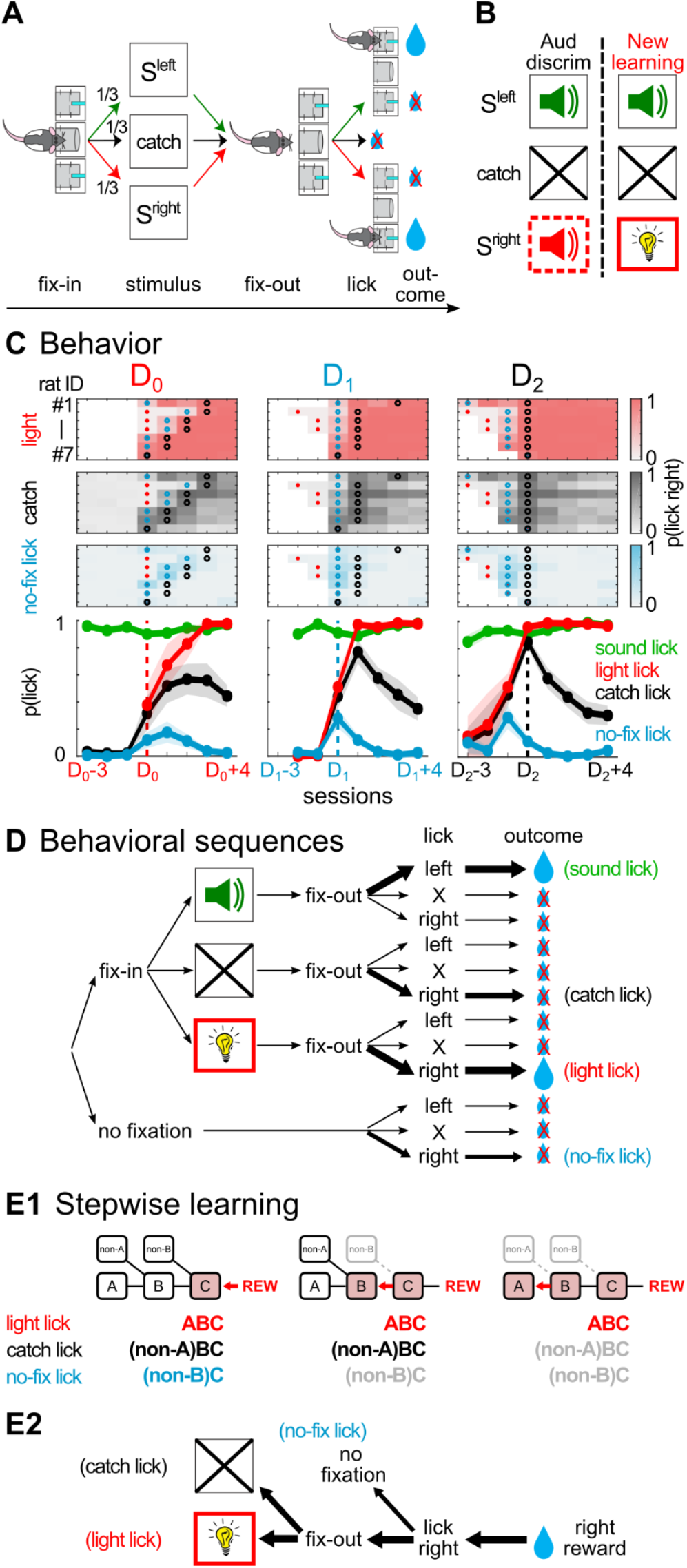
New associative learning led to sequential refinements of reward-seeking behaviors. **A**, Behavioral task. Rats entered the fixation port to initiate each trial, where they encountered three trial types (S^left^; S^right^; catch) with equal probabilities that respectively indicated water reward in the left or right port, or no reward in the case of catch trials (no stimulus). **B**, Rats initially learned an auditory discrimination task (old association phase). At the new learning phase, S^right^ was switched from the sound to a new house light, while other elements of the task remained the same. **C**, The proportion of three types of reward-seeking behaviors toward the right reward port (light licks, catch licks, no-fixation licks) across sessions during new learning (N=7 rats). No fixation licks (cyan) refers to trials in which rats failed to first enter the fixation port before licking in the right reward port. Sessions were respectively aligned, in each column, at the D_0_, D_1_ or D_2_ session of each animal. D_0_ refers to the first new learning session with the new light stimulus; D_1_ refers to the session when animals began to respond correctly in the new light trial; D_2_ refers to the session when catch licks peaked. The D_0_, D_1_ and D_2_ sessions in each animal were indicated by red, cyan, and black circles, respectively. Each row in top panels depicts behavior performance in one animal (#1-7). In the middle and right panels, only sessions in the new learning phase (starting from the D_0_ session) were plotted. The emergence, as well as the following sequential elimination, of no-fixation licks and catch licks resembled the sequential refinement of reward-seeking behaviors under the stepwise learning strategy. **D**, Behavioral sequences that animals might experience. Out of all possible sequences, only three types of rightward licking behaviors were consistently observed during new learning. **E1**, The pattern of the three types of rightward licking behaviors rearranged as the discrete learning steps as in Figure 1B. **E2**, The corresponding behavioral events that constituted the three types of rightward licking behaviors, arranged as in Figure 1B.

To understand how the repertoire of behavioral sequences evolved during learning, we examined all possible behavioral sequences that the animal might experience (Figure 2D). This approach allowed us to identify non-rewarded behavioral sequences that were not associated with specific trial types but were, nonetheless, highly relevant for the learning process. We identified three types of behavioral sequences whose frequencies consistently increased during the new learning phase. These three types of behavioral sequences included the rewarded licking behavior in the new light trials (light licks), as well as two types of non-rewarded behavioral sequences: catch licks and no-fixation licks (Figure 2C). Catch licks refers to licking responses toward the right reward port in catch trials when no sensory stimulus was presented. No-fixation licks refers to the situation in which rats directly licked at the right reward port without first entering the center fixation port. All three behavioral sequences shared the common feature of licking the right reward port (lick-right).

Both no-fixation licks and catch licks were largely absent before the new learning phase, and emerged and peaked during early sessions of new learning, before subsequently diminished in later sessions (Figure 2C). No-fixation licks occurred most frequently at the D_1_ session, while catch licks peaked in the same session (N=1/7) or later in most animals (N=6/7). We will denote the peak of catch licks as the D_2_ session in each animal. The consistent temporal order of the D_1_ and D_2_ sessions within each animal allowed us to identify similar learning stages across animals despite their individual differences in learning dynamics.

The temporal dynamics of the three types of rightward licks during new learning (Figure 2C) showed that non-rewarded behaviors were sequentially eliminated while rewarded behaviors were preserved. This temporal dynamics resembled the pattern of sequential refinement of behavioral sequences predicted by the stepwise learning strategy (Figure 2E), and likely corresponded to the discrete steps in the underlying learning process. To further test whether such patterns of sequential refinements represent a general feature of behavioral sequence learning, we trained a separate cohort of animals and observed similar behavioral signatures regardless of the sensory modality of the stimulus or the laterality of the new learning side (Figure S1). These observations support the idea that animals do use the stepwise strategy to learn behavioral sequences. In the following analyses, we tested additional predictions of the stepwise learning strategy at both behavioral and neurophysiological levels.

### Initial learning was characterized by the rapid emergence of reward-seeking behaviors and corresponding increases in BF activities

To validate the behavioral observations and understand the underlying neural dynamics, we recorded BF neuronal activity throughout the learning process (Figure 3A) and used the consistent S^left^ sound as the control stimulus to identify stable populations of BF bursting neurons (Figure 3B, Figure S2). A total of 1453 BF single units were recorded over 45 sessions (N=7 rats), of which 70% (1013/1453) were classified as BF bursting neurons based on their stereotypical phasic response to the S^left^ sound (22.5±7.3 neurons per session, mean±std) (Figure 3B). The population response of BF bursting neurons were highly consistent across animals (Figure 3C), and remained remarkably stable in S^left^ trials throughout the learning process (Figure 3D). The inclusion of S^left^ trials therefore allowed us to record from stable and representative populations of BF bursting neurons throughout the learning process, and to investigate how their activities dynamically evolved in other trial types during learning at single trial resolution (Figure 3E).

**Figure 3.**
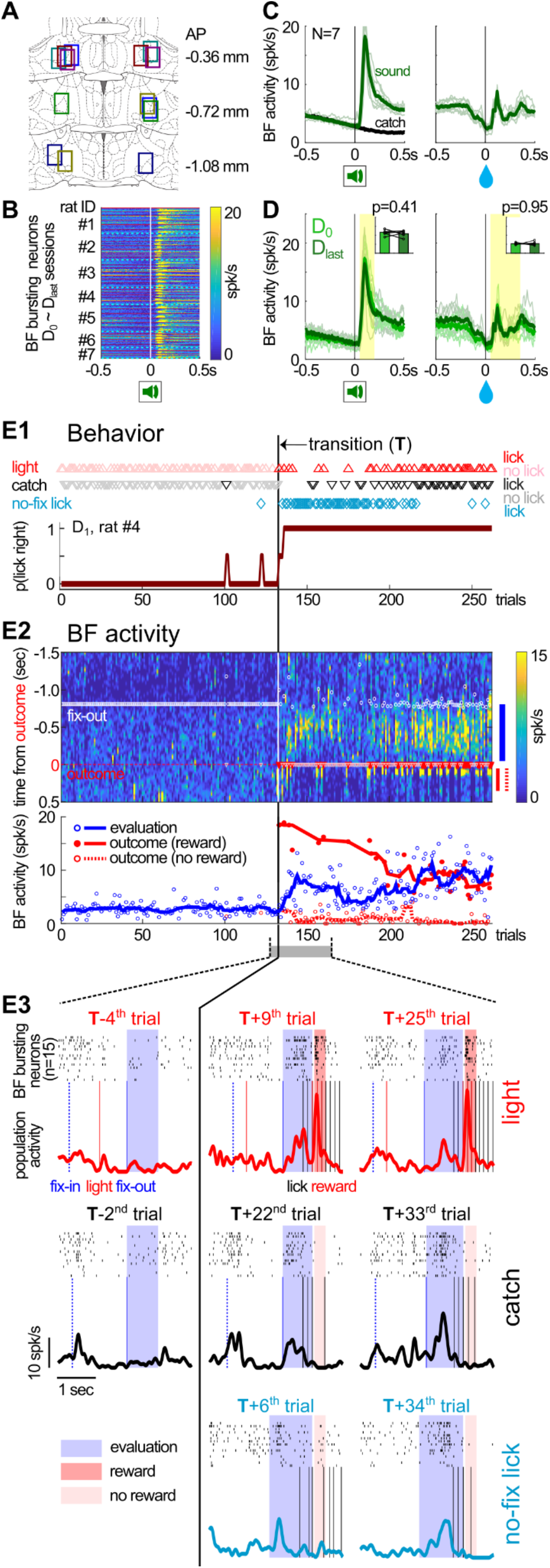
Abrupt transition in reward-seeking behaviors corresponded to increased neuronal activity in the BF during initial learning. **A**, Locations of electrode bundles targeting bilateral BF (N=7) in coronal sections of the rat brain (coordinates relative to Bregma). Different colors correspond to different animals. **B**, Response of individual BF bursting neurons (n=1013) to the S^left^ sound during new learning sessions (N=45 sessions; separated by thin red lines) in each animal (N=7 rats; separated by cyan dotted lines). BF bursting neurons showed robust and consistent phasic responses to the S^left^ sound throughout the learning process. **C**, Average BF bursting neuron responses to S^left^ sound onset and the associated reward delivery. BF activities in catch trials were plotted for comparison. Responses from individual animals (thin lines) were similar. **D**, The activity of BF bursting neurons remained stable between the first (D_0_) and last (D_last_) recording session. Average activities in the yellow shaded intervals were similar between these two sessions (inset). Thin lines indicate BF activity in individual animals. **E**, Behavioral and BF neuronal dynamics in the D_1_ session from a representative animal (rat #4). **E1**, The emergence of three types of rightward licks after the transition point (top), and their combined rightward licking probability across trial types (bottom). The transition point (T) marked an abrupt transition in the pattern reward-seeking behavior that went from no licking to 100% licking. **E2**, Top, population activities of BF bursting neurons (color-coded) in the same trials (X-axis) as shown in E1. Y-axis indicates time in each trial, with time zero aligned at the trial outcome. No lick trials before the transition were aligned instead at the time of fixation port exit (fix-out, white circle) such that the median timing of fix-outs in lick and no lick trials were equivalent. The blue and red lines to the right of the panel indicate the time windows for calculating evaluation and outcome responses, respectively. Bottom, BF evaluation responses (blue) and outcome (red) responses across trials. Outcome responses were plotted separately for rewarded (solid red) and non-rewarded (dashed red) licks. Circles indicate BF activities in single trials and lines indicate their respective trends (moving medians). **E3**, Examples of single trial BF activities from the three types of rightward licks taken around the transition point. Each panel showed the spike rasters of BF bursting neurons in this session (n=15) (top), along with the population activity trace and relevant behavioral events (bottom). Shaded intervals indicate the time windows corresponding to evaluation responses (blue) and outcome responses (red) shown in E2. Notice that BF bursting neurons in the same session showed highly similar activity patterns, and that the BF evaluation response rapidly emerged in all three types of rightward licks after the transition point.

We first applied this approach to understand the behavioral and neural dynamics in the D_1_ session because all three types of rightward licks emerged in this session (Figure 2C, middle panel). Detailed analysis of behavioral responses from a representative session (Figure 3E) revealed that the three types of rightward licking behaviors emerged abruptly after a transition point (see Methods for definition). Rightward licking behaviors were mostly absent before the transition point, and rapidly switched to almost 100% licking after the transition point. This pattern was consistently observed across all animals (Figure 4A1).

**Figure 4.**
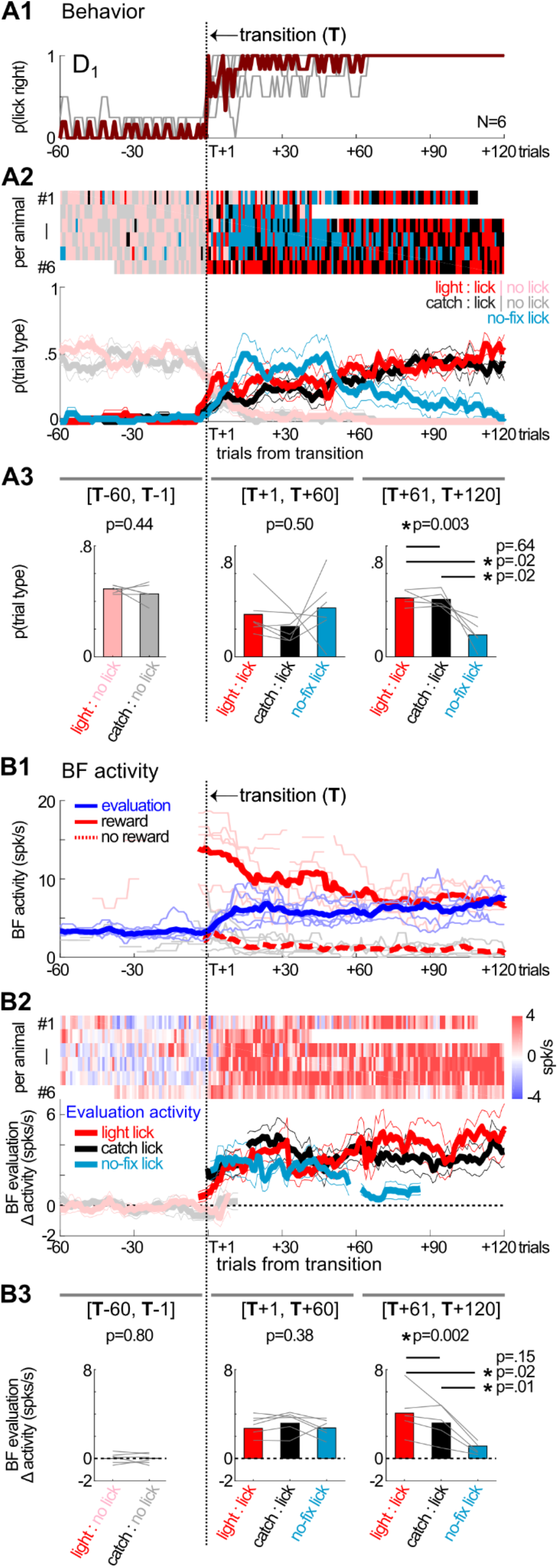
Rapid emergence and subsequent refinement of reward-seeking behaviors and BF activity. **A**, The pattern of rightward licking behaviors aligned at the transition point in the D_1_ session of each animal (N=6). One animal (#7) with accelerated learning dynamics, in which its D_1_ and D_2_ occurred in the same session, was excluded from this analysis. The behavioral response patterns in three trial types (light trials, catch trials and no-fixation licks) were either pooled together within each animal (**A1**), or plotted separately (mean ± s.e.m.) (**A2**). While all three types of rightward licking emerged immediately after the transition, no-fixation licks subsequently decreased after 60 trials (**A3**). **B**, Corresponding changes in BF activity aligned at the transition point in the D_1_ session of each animal (N=6). BF evaluation responses and outcome responses in the three trial types were pooled within each animal (**B1**), as in the example in Figure 3E2. BF evaluation responses were further plotted separately for each trial type, relative to their respective baseline firing rates (mean ± s.e.m.) (**B2**). BF evaluation responses increased similarly in the three types of rightward licking immediately after the transition, and subsequently decreased in no-fixation licks after 60 trials (**B3**). Thin lines in B1 indicate the trend (10-trial moving median) of BF activities from individual animals.

At the neuronal level, there was a corresponding increase in the activity of BF bursting neurons that rapidly emerged after the transition in reward-seeking behaviors (Figure 3E, 4B1). This increase in BF activity was most prominent in the epoch before the trial outcome as animals approached the reward port (Figure 3E2). In contrast, in trials before the transition point, BF bursting neurons did not show similar activity increases in the corresponding time window after exiting the fixation port (Figure 3E, 4B1). We will refer to the BF activity in this window as the BF evaluation response (see Methods for definition) because it reflected animals’ internal evaluation when no sensory stimuli were presented during this epoch.

### The emergence and subsequent elimination of non-rewarded behaviors were mirrored by changes in BF activity

The stepwise learning model predicted that, as animals learned about the lick-right action as the first reward predictor, animals would initially engage in all three types of rightward licking behaviors because they all contained this reward predictor (Figure 2E). We therefore examined the respective prevalence of the three types of rightward licking behaviors after the transition point in the D_1_ session (Figure 4A). All three types of rightward licking behaviors emerged immediately after the transition point. Moreover, after about 60 trials, no-fixation licks began to decline and occurred less frequently than light licks and catch licks (Figure 4A2-3).

At the neuronal level, the corresponding prediction was that the lick-right action would be associated with increases in BF activities during the first learning step, which should be similarly present in all three types of rightward licking behaviors. Indeed, BF evaluation responses quickly increased after the transition point in all three types of rightward licking behaviors (Figure 4B). The amplitudes of BF evaluation responses were similar between the three types of rightward licking behaviors within the first 60 trials after the transition. Subsequently, the BF evaluation response in no-fixation licks declined in the next 60 trials, relative to the other two types of rightward licking behaviors (Figure 4B2-3).

These results support that the first step of the stepwise learning process corresponded to the first 60 trials after the transition point in the D_1_ session. All three types of rightward licking behaviors were present during this step of learning. BF activity also increased to similar levels in all three types of behavioral sequences whenever animals approached and licked the right reward port, regardless of whether they had exited from the center fixation port.

These results further suggest that the second step of learning started at roughly 60 trials after the transition, at which point animals learned about the second reward predictor: exiting the fixation port (Figure 2E). As a result, the no-fixation lick sequence was no longer compatible with the learned reward predictors, which resulted in diminished BF evaluation responses and the elimination of this behavior from the behavioral repertoire. The other two types of rightward licks, light licks and catch licks, remained compatible and maintained high levels of BF evaluation responses.

### BF neurons did not respond to the new light stimulus during initial learning

A further prediction of the stepwise learning model was that, at both the first and the second steps of learning, animals had not learned to use the new light stimulus as a reward predictor, and therefore the new light stimulus would not elicit responses in BF bursting neurons during these steps (Figure 2E).

We tested this prediction by comparing BF activities between light and catch trials in the D_1_ session (Figure 5). Indeed, BF activities in light trials were highly similar to those in catch trials (in the absence of the light stimulus), in the epochs after stimulus onset as well as before exiting the fixation port. This was true regardless of whether animals subsequently licked at the right reward port. This observation confirmed the prediction that the new light stimulus did not activate BF bursting neurons in the D_1_ session, despite the near-perfect behavioral performance in light trials after the behavior transition point.

**Figure 5.**
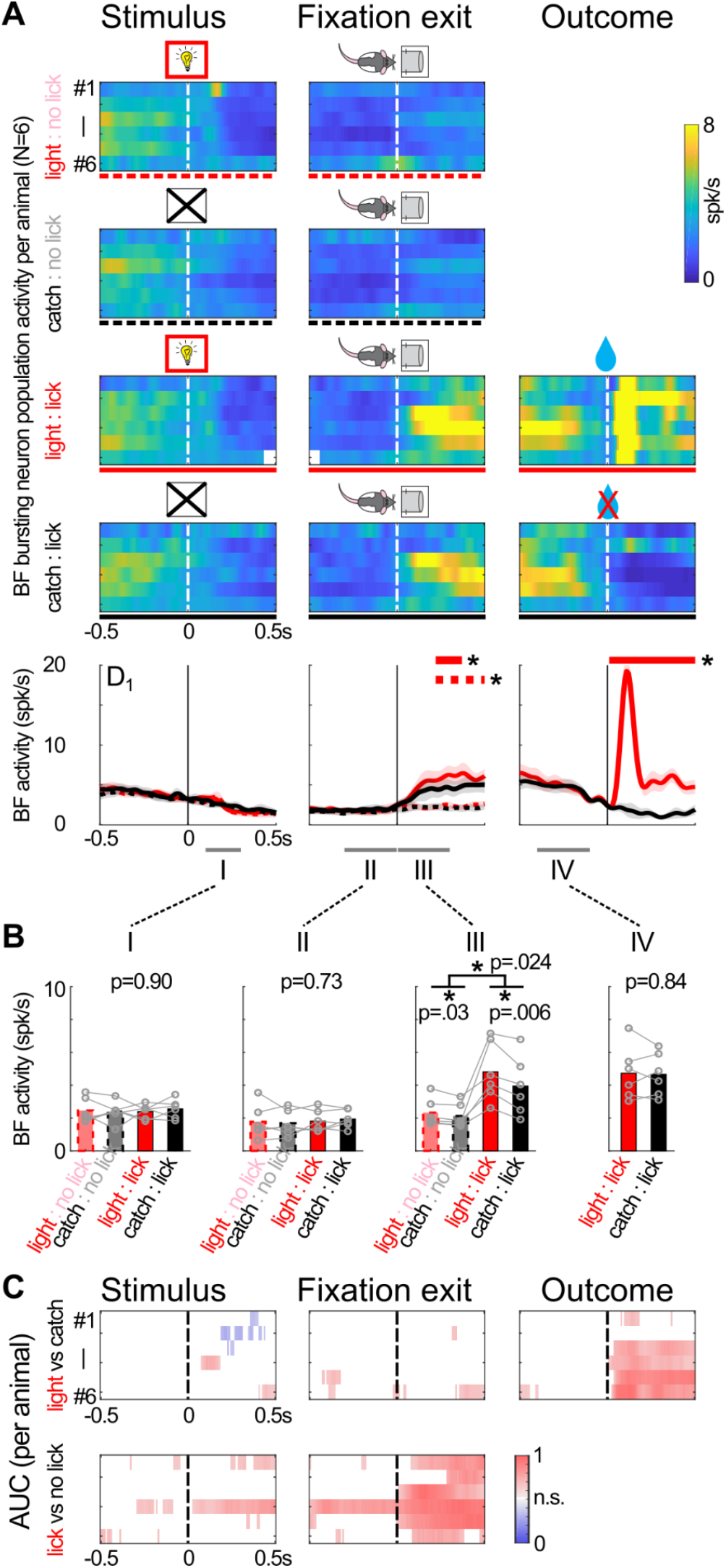
BF neurons did not respond to the new light stimulus during initial learning. **A**, Population activities of BF bursting neurons in light and catch trials in the D_1_ session. BF activities from individual animals (N=6) (top) and group averages (mean ± s.e.m.) (bottom) were plotted separately based on trial type (light or catch) and behavioral response (licking at the right reward port or not), and aligned at three key behavioral events in each trial: stimulus onset, fixation port exit, and trial outcome. The solid and dashed lines below the top panels indicate the line symbol used in the bottom panels. Horizontal lines in bottom panels indicate significant differences in BF activities (p<0.01, 3 consecutive bins) between light lick and catch lick trials (solid red), or between lick and no lick trials (dashed red). **B**, Comparison of BF activities between the four trial combinations during four time windows (I-IV). There was no difference in BF activities before exiting the fixation port (I & II). In contrast, BF activities increased in both light and catch lick trials after exiting from the fixation port (III & IV). BF activities in light lick and catch lick trials were similar before the trial outcome (IV). **C**, Comparison of BF activities between light and catch trials (top) or between lick and no lick trials (bottom) using sliding window ROC analysis, aligned at the three behavioral events. Only significant (p<0.001) area-under-curve (AUC) values were shown. BF activities could not reliably discriminate between light and catch trials before the receipt of reward. In contrast, BF activities could reliably discriminate between lick and no lick trials after exiting the fixation port.

On the other hand, in the epoch after exiting the fixation port, there were similar increases in BF activity when animals licked at the right reward port in both light and catch trials (Figure 5). This increase in BF activity corresponded to the BF evaluation response described earlier (Figure 4B), which reliably distinguished between lick and no lick trials, but not between light and catch trials (Figure 5C). These observations support the idea that light and catch trials were treated as the same in the D_1_ session, and that the light stimulus has not been learned as a reward predictor at this stage of learning.

### BF responses to light onset emerged later when the light stimulus was used to guide reward-seeking behavior

When did the third step of the stepwise learning process take place? Since the third step was when animals learned to use the light stimulus as a reward predictor to guide reward-seeking behaviors, it should correspond to the time point when the behavioral performance in light and catch trials began to diverge. We noted that the behavioral pattern in light and catch trials were highly similar prior to the D_2_ session (Figure 2C), and the similarity was best illustrated in the fine behavioral and neuronal dynamics within the D_1_ session (Figure 4). In contrast, during the D_2_ session, the behavioral pattern in light and catch trials began to show a small but significant difference (Figure 6A). This pattern suggests that the D_2_ session might be when the third step of learning took place.

**Figure 6.**
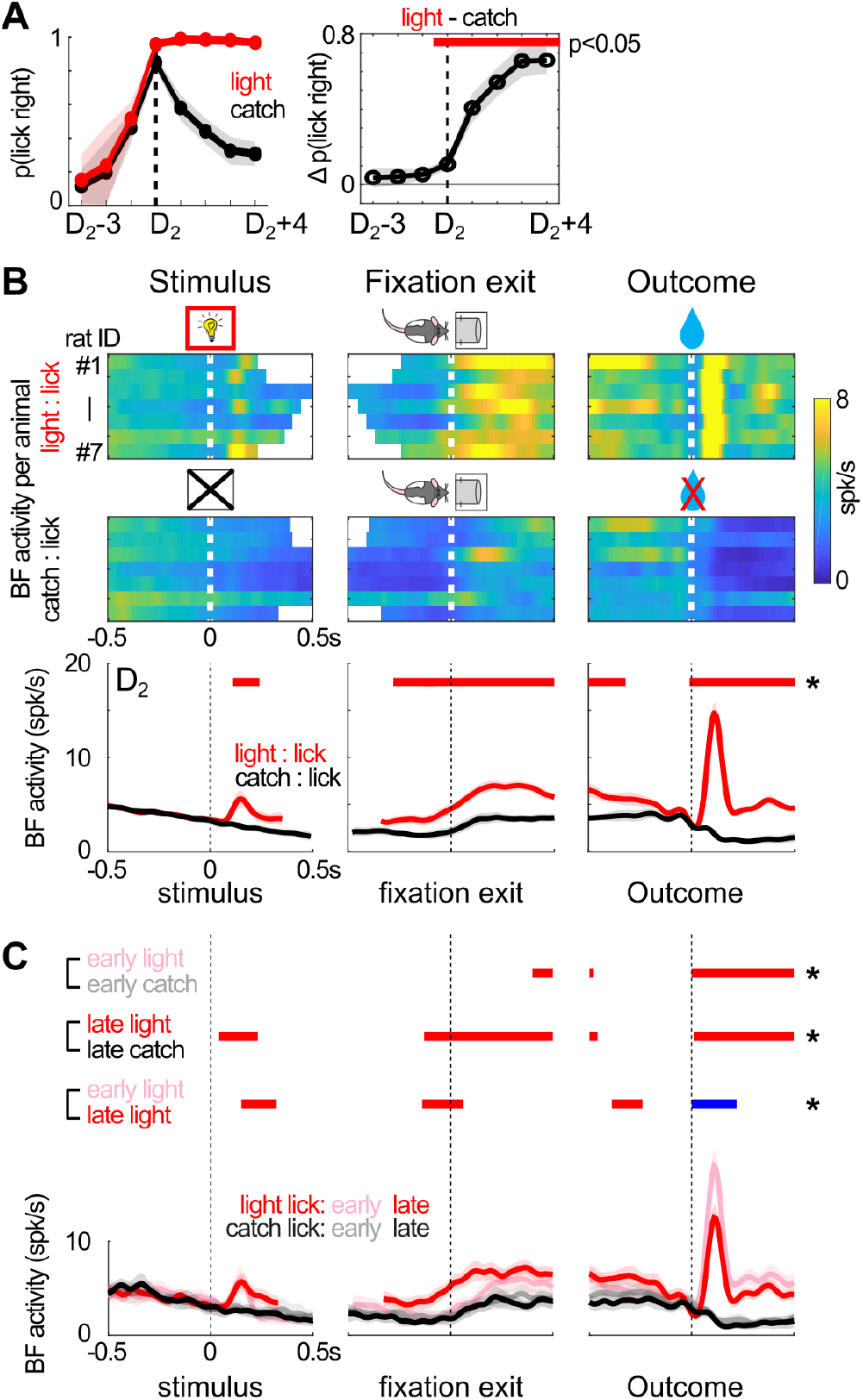
BF responses to light onset emerged later when light was used to guide reward-seeking behavior. **A**, The probability of rightward lick in light and catch trials (left), aligned to the D_2_ session of each animal (N=7), began to differ significantly at the D_2_ session (p<0.05) (right). **B**, Population activities of BF bursting neurons in light lick and catch lick trials in the D_2_ session for individual animals (top) (N=7) and their group average (mean ± s.e.m.) (bottom). BF activities in light lick trials were significantly higher in epochs before exiting the fixation port. Significant differences in BF activities (p<0.01, 3 consecutive bins) were indicated by horizontal red lines. BF activities were truncated at the median RT of the respective sessions (see Methods for details). **C**, BF activities (mean ± s.e.m.) in light lick and catch lick trials at the first 20 trials (early) or the last 20 trials (late) of the D_2_ session. At the beginning of the D_2_ session, BF activities between light lick and catch lick trials were similar in the interval between stimulus onset and fixation port exit. Differences in BF activities in this interval became significant by the end of the D_2_ session. Similar differences in this interval were observed when comparing BF activities between the early and late light lick trials. Significant excitation (red) or inhibition (blue) in BF activities (p<0.01, 3 consecutive bins) were indicated by horizontal lines.

The corresponding prediction at the neuronal level was that BF bursting neurons should begin to respond to the light stimulus in the D_2_ session. We tested this prediction by comparing BF activities between light and catch lick trials in the D_2_ session, and found significant differences in all epochs, including the presence of a phasic response to the light onset (Figure 6B). This pattern was distinct from the pattern of BF activities in the D_1_ session, when there was no difference in the epochs before exiting the fixation port (Figure 5). We further investigated whether this difference in BF activities between light and catch lick trials emerged within the D_2_ session. To test this idea, we compared BF activities at the beginning (first 20 trials) and the end (last 20 trials) of this session (Figure 6C). We found that, at the beginning of the D_2_ session, BF activities patterns were similar to the pattern in the D_1_ session (Figure 5A), showing no difference between light and catch trials before exiting the fixation port. However, by the end of the D_2_ session, BF responses to the light stimulus had developed in the epochs before exiting the fixation port (Figure 6C). These results further support the idea that the third step of learning took place in the D_2_ session, which was when animals first learned about the light as a reward predictor.

### Stronger BF responses to the new light reflected better learning and faster decisions

After the third step of learning took place in the D_2_ session, did the learning about the new light plateaued or did the learning continue to progress? At the neuronal level, BF phasic responses to the new light continued to grow stronger after the D_2_ session (Figure 7A). At first glance, the continual increases in BF responses to the new light stimulus did not match with the hit rates in light trials, which had already plateaued in the D_2_ session (Figure 2C). However, as we have shown earlier (Figures 4 & 5), hit rates in light trials could be a poor index of learning about the new light stimulus because light licks in the early learning sessions were not driven by the light stimulus but by later events in the behavioral sequence (exiting fixation and lick-right) as reward predictors. Those behavioral events enabled rightward licking responses in the absence of the light stimulus (catch licks and no-fixation licks).

**Figure 7.**
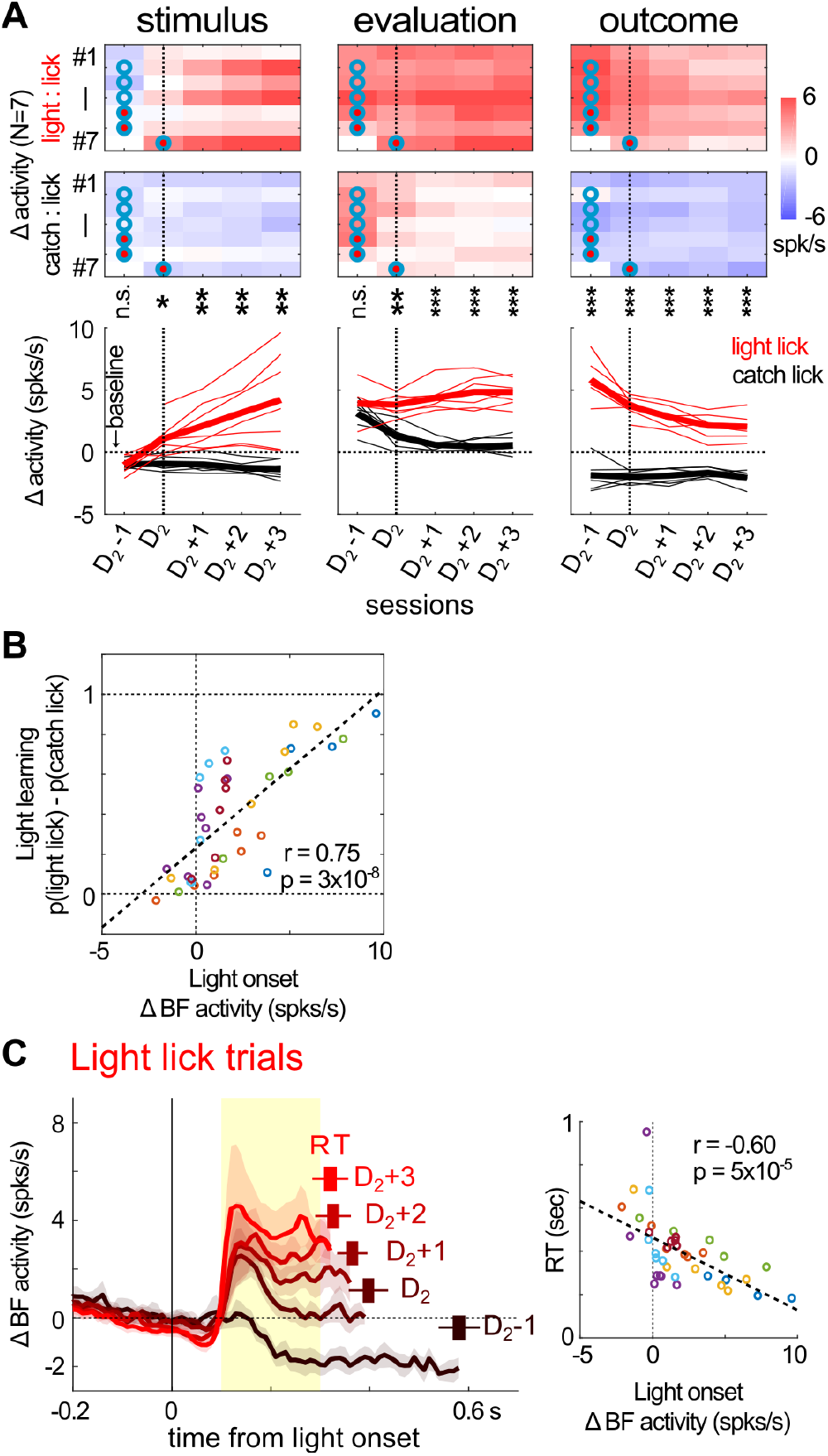
Increased BF responses to light onset predicted the extent of light learning and faster reaction times. **A**, The evolution of BF activities in light and catch lick trials, aligned at each animal’s D_2_ session, calculated relative to their respective baseline firing rates. Significant differences in BF activities between the two trial types were indicated (*p<0.05; **p<0.01; ***p<0.001). BF activities in the stimulus onset and evaluation epochs diverged in the D_2_ session and their differences grew larger afterwards. Also notice that BF responses to the reward in light lick trials decreased over sessions. **B**, The average amplitude of BF responses to light onset in light lick trials was positively correlated with the behavioral index of light learning, i.e. the difference in the probability of rightward licks between light and catch trials. BF activities were calculated relative to their respective baseline firing rates in each session. Each circle indicates one session and different colors correspond to different animals. **C**, Average responses of BF bursting neurons (mean ± s.e.m.) to light onset in light lick trials (left). BF activities, relative to their respective baseline firing rates, were truncated at the median RT of the respective sessions. The RTs (mean ± s.e.m.) of the corresponding sessions were shown above each trace. Stronger BF responses to light onset in the [0.1, 0.3]s window (yellow shaded interval, left panel) were correlated with faster RTs in individual sessions (right). Each circle indicates one session and different colors correspond to different animals, same as in **B**.

Instead of the hit rate in light trials, a better behavioral index for the learning about the light stimulus would be the difference in the levels of behavioral performance between light and catch trials. This behavioral index accounted for the contributions from the later events in the behavioral sequence that were shared between light and catch licks, and isolated the contribution of the light stimulus to the rightward licking behavior. We found that this index of light learning was strongly correlated with the amplitude of BF phasic responses to the light stimulus in individual sessions (Figure 7B). This observation therefore supports the idea that the learning about the light stimulus continued to grow stronger after the D_2_ session.

Another dimension of the learning about the light stimulus was the change in animals’ decision speeds, measured by reaction times (RTs). Previous studies have shown that stronger BF bursting responses are quantitatively coupled with, and causally lead to, faster RTs (*21, 22*). Such observations predicted that there would be corresponding decreases in RTs toward the light stimulus after the D_2_ session. In support of this prediction, we found that stronger phasic bursting responses to light onset were coupled with faster RTs in individual sessions (Figure 7C). Together, these observations support the idea that the learning about the new light was reflected by the amplitude of BF phasic response to the light stimulus.

In addition, BF activities not only reflected the learning about the light stimulus, but also predicted reward-seeking behaviors in the absence of the light stimulus. For example, in catch trials, stronger BF activities after exiting the fixation port predicted rightward licking behavior and discriminated lick trials from no lick trials (Figure S3). Moreover, increased BF activities before the start of licking predicted longer durations of licking in catch lick trials when no reward was delivered (Figure S3). These observations provided additional support for the idea that the activity of BF bursting neurons promoted reward-seeking behaviors.

### Reward expectations negatively modulated BF responses to trial outcomes

A final validation of the learning dynamics came from how BF responses to the trial outcome were modulated by reward expectations and BF activities in earlier epochs. Previous studies have found that the response of BF bursting neurons to the reward was negatively modulated by reward expectation (*21, 22, 27, 28*). Such properties would predict that, in the current experiment, BF responses to the reward should decrease throughout the stepwise learning process. Indeed, BF responses to the right reward decreased over trials in the D_1_ session (Figures 3E2 & 4B1), and continued to decrease over subsequent sessions (Figure 7A). These results support that animals learned to better predict the rewarding outcome across different steps of the learning process.

We further investigated whether the response of BF bursting neurons to trial outcomes were negatively correlated with BF responses in earlier epochs. Such a negative correlation is a hallmark feature of reward prediction error encoding (*29*) and has been previously reported in BF bursting neurons (*21, 22*). Indeed, at the per session level, we found that the amplitude of BF responses to the reward in light trials was strongly and negatively correlated with the amplitude of BF phasic bursting response to the light onset (Figure 8A). Moreover, in pre-D_2_ sessions where BF responses to the new light stimulus had yet to develop, we found that BF responses to the trial outcome were negatively correlated with BF evaluation responses in the same trial (Figure 8B). This negative correlation at the single trial level was observed not only when the reward was delivered (light licks) but also when the reward was absent (no-fixation licks and catch licks) (Figure 8B, 8C). The fact that these patterns were observed in catch licks and non-fixation licks supports that animals were expecting to receive reward in those trials, and the extent of reward expectation was similarly reflected in BF evaluation responses and negatively modulated BF responses to the trial outcome. Together, these observations support that BF bursting neurons encoded reward prediction error, and that BF activities in earlier epochs (stimulus onset or evaluation window) reflected animals’ reward expectations.

**Figure 8.**
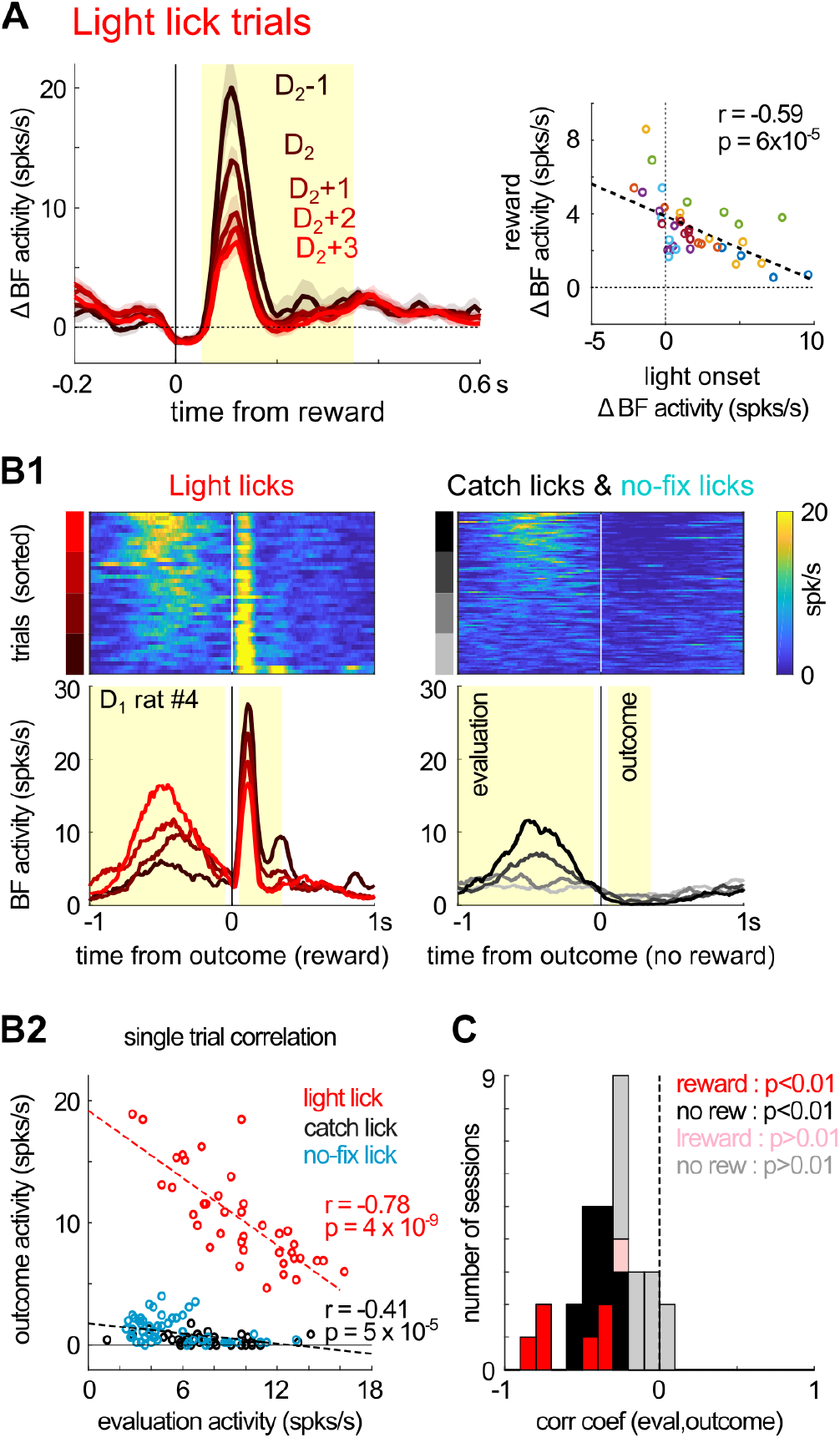
BF responses to trial outcome were negatively correlated with BF activities in earlier epochs. **A**, Average responses of BF bursting neurons (mean ± s.e.m.) to the reward in light lick trials, relative to their respective baseline firing rates, plotted separately for the five sessions relative to the D_2_ session (left). BF responses to the reward (yellow shaded interval, left panel) were negatively correlated with BF responses to light onset (Figure 7C) in individual sessions (right). Each circle indicates one session and different colors correspond to different animals. **B**, Negative correlation of BF activities between evaluation and outcome responses in single trials. **B1**, Single trial BF activities in an example D_1_ session, aligned at the trial outcome. Single trial BF responses were plotted separately for rewarded licks (light licks) and non-rewarded licks (catch licks and no-fixation licks). Yellow shaded intervals indicate time windows for calculating evaluation and outcome responses. Trials were sorted by the amplitude of evaluation responses (top). Average BF activities from the four quartiles of trials were plotted separately (bottom). **B2**, Negative correlation between single trial BF evaluation responses and outcome responses in this session. Each dot represents one trial. Catch licks and no-fixation licks were pooled together to calculate the correlation in non-rewarded licks. **C**, Histogram of correlation coefficients between evaluation and outcome responses from individual sessions. Results for rewarded lick trials (light licks) were calculated from pre-D_2_ sessions when BF responses to the light stimulus had not developed (N=7), and for non-rewarded lick trials from sessions with at least 50 trials of catch and no-fixation licks combined (N=25). Most sessions showed significant negative correlations.

## Discussion

Results from the current study support that animals used a stepwise strategy to learn behavioral sequences that contain both sensory stimuli and their own actions. Behavioral events were learned sequentially, starting from the event closest to the reward and sequentially expanded to earlier events (Figure 1). The behavioral signature of this stepwise learning process was the sequential refinement of rightward licking behaviors, in which non-rewarded licking behaviors (no-fixation licks and catch licks) were sequentially eliminated while the rewarded behavior (light licks) was preserved (Figure 2). Learning about each behavioral event as a new reward predictor was accompanied by the emergence of BF responses toward that event, which conveyed animals’ reward prediction toward that event. Increased BF activities first emerged in the epoch before animals entered the reward port (Figures 3E & 4B), while responses to the earlier event (light stimulus) developed later (Figures 5 & 6). The evolution of BF activities mirrored the behavioral response patterns, which was initially increased in all three types of rightward licks (Figures 3E & 4B) and subsequently decreased in non-rewarded licks as those behaviors were eliminated (Figures 4B & 7A). Throughout the learning process, the activity of BF bursting neurons encoded reward prediction error signals (Figure 8) and their increased activities consistently predicted reward-seeking behaviors and faster reaction times (Figure 7).

These results therefore identified the behavioral and neurophysiological signatures of the stepwise learning strategy when animals learned behavioral sequences.

### Stepwise learning of behavioral sequences

The current study focused on the learning of behavioral sequences that contain both sensory stimuli and actions, which reflect behavioral contexts that are commonly encountered both in experimental and natural settings. The learning dynamics that we described in this study cannot be easily accounted for using either Pavlovian (*1, 2*) or instrumental (*3, 4*) conditioning alone. In this regard, stepwise learning provides a new framework to understand the learning of behavioral sequences.

Theoretically, long behavioral sequences are difficult to learn because the number of sequence permutations grows exponentially as a function of the sequence length. However, modeling studies have suggested that such learning can be greatly accelerated using the stepwise strategy, which reduces the number of sequence permutations to a linear function of the sequence length (*18*). The current study provides behavioral and neurophysiological evidence to support that animals indeed adopt the stepwise learning strategy to learn behavioral sequences. At each step of learning, animals explored the subset of behavioral sequences that shared common sequence elements learned in previous steps (Figures 1 & 2). Such explorations allowed animals to distinguish those sequences, which were initially indistinguishable at the beginning of that learning step, and selectively eliminate subsets of non-rewarded sequences. From this perspective, the stepwise learning strategy offers an intuitive explaination of why animals committed certain types of ‘errors’ (non-rewarded licks), and suggests that those behaviors in fact represented genuine reward-seeking efforts at earlier stages of learning. By learning the associative relationship one step at a time, the stepwise learning strategy likely minimized the cognitive burden of the animal during the learning process and enabled the efficient learning of complex sequences.

Our results suggest that stepwise learning is likely a general strategy for learning behavioral sequences because its behavioral and neural signatures were also observed in other learning settings. In a separate cohort of animals, we showed that the behavioral signatures of sequential refinements were similarly observed when the sensory modalities of the stimulus were reversed, and regardless of the laterality of the new learning side (Figure S1). Moreover, even during the learning of the most simple behavioral sequence containing only two elements (stimulus-action), similar behavioral and neural signatures were also observed (Figure S4).

It is important to note that, while these results support that stepwise learning is a widely-used strategy for learning behavioral sequences, they do not preclude other possible learning strategies. In particular, if animals had previously encountered similar behavioral events, such experiences could be generalized to the new learning context, and allow animals to use a forward chaining strategy to learn behavioral sequences, instead of the backward chaining order in the stepwise learning strategy (*18*). For example, if animals had previously learned that light stimuli could predict reward, they could generalize that experience to the current learning task and view the new light stimulus as a potential reward predictor. Such generalizations would lead animals to engage in reward-seeking behaviors in light trials and quickly discover the light-reward association, bypassing the learning of behavioral events later in the sequence. Such strategies perhaps underlie the accelerated learning dynamics that we observed in a subset of animal (animal #7, Figure 2C), which also featured the strongest BF responses to the light stimulus during the initial encounter of the new light (Figure 6B).

### Temporal dynamics of the new learning process

Our results revealed that the true temporal dynamics about the learning of the new stimulus can be very different from the dynamics of behavioral performance levels in the new stimulus trial. We found that, despite the near-perfect behavioral performance in light trials that began after the transition point in the D_1_ session (Figures 2-4), the learning of the new light stimulus did not begin until the D_2_ session (Figures 5-6). Moreover, the learning of the new light stimulus continued to grow after the D_2_ session despite the plateaued behavioral performance in light trials (Figures 6-7). During early stages of learning, while the light stimulus was clearly perceptible and had been consistently paired with reward over many trials, the reward-seeking behavior in light trials was not driven by the light stimulus but by later behavioral events in the sequence (fixation port exit and lick-right). The critical factors that allowed us to reach this conclusion were the analysis of non-rewarded licking responses and the inclusion of catch trials in our task design. If not for these factors, we would incorrectly conclude that the learning about the new light stimulus occurred much earlier.

The discrepancy between behavioral performance in light trials and the learning about the light stimulus highlights the potential mismatch in which salient physical events (such as the light stimulus) are not always automatically used by animals to predict reward, especially during the early stages of learning. The light stimulus was only incorporated as a reward predictor in later stages of the learning process, despite its continued presence from the beginning of the new learning. This observation indicates that, even in the absence of changes in reward contingencies in the environment, the learning process can lead to the addition of new reward predictors. Such structural revisions of the internal reward prediction model pose a fundamental challenge to theories and models of learning that assume a static set of reward predictors. Using models with incorrect reward predictors will lead to incorrect interpretations, regardless of how well the model fits the behavioral performance.

### Neural correlate of stepwise learning in the activity of BF bursting neurons

The current study extends our understanding about the roles of BF bursting neurons in the encoding of reward prediction error. Previous studies have demonstrated that BF responses to rewards are negatively modulated by reward expectation (*21, 22, 27*) and support the idea that BF neurons encode a reward prediction error signal (*27, 28*). The current study further extends this idea and shows that BF bursting neurons similarly encode reward prediction error in the context of new learning, and such encoding is robust even at the single trial level (Figure 8). This robust encoding of reward prediction error by BF bursting neurons quantitatively conveys the amount of reward prediction associated with each behavioral event at each learning step. As a result, the stepwise learning process was mirrored by the activity of BF bursting neurons, which provides a neural correlate of the stepwise learning process that we were able to track throughout the learning process in single trials.

Previous studies have also established that BF bursting neurons serve as a bidirectional gain modulation mechanism for reward-seeking behaviors, where increased BF activities promote faster reaction times (*21, 22*) while the inhibition of BF activities leads to rapid behavioral stopping (*23*). Manipulations of BF activities using electrical stimulation further suggest that BF bursting neurons likely play a causal role that can modulate decision speed when their activities are increased (*22*) or inhibited (*23*). The current study extends this idea to the context of new learning, by showing that increased BF activities were tightly coupled with reward-seeking behaviors at multiple levels throughout the learning process (Figures 3-7), and quantitatively predicted faster reaction times (Figure 7) and longer licking durations (Figure S3). The engagement in reward-seeking behaviors was particularly important in the context of learning behavioral sequences, because such explorations were essential for discovering the relationship between earlier events in the behavioral sequence and the rewarding outcome. Taken together, BF bursting neurons may serve the role of transforming the encoding of reward-prediction error into promoting reward-seeking behaviors during the new learning process.

Finally, several studies have suggested that BF bursting neurons are likely a subset of GABAergic neurons (*21, 24, 30*), but their specific cellular marker(s) remains to be determined. Such marker information will be needed to conduct selective manipulation experiments that specifically target BF bursting neurons to directly test their causal role in the learning process.

## Methods

### Ethics Statement

All experimental procedures were conducted in accordance with the National Institutes of Health (NIH) Guide for Care and Use of Laboratory Animals and approved by the National Institute on Aging (NIA) Animal Care and Use Committee and by the Institutional Animal Care and Use Committee at the National Yang Ming Chiao Tung University, Taiwan (NYCU).

### Subjects

Seven male Long Evans rats (Charles River, NC), aged 3–6 months and weighing 300–400 grams were used for the recording experiment. Rats were housed in a 12/12 day/night cycle and were provided with 10 to 12 dry pellets per day and unrestricted access to water. Rats were trained in daily sessions lasting 60-90 minutes. A separate cohort of eight male Long Evans rats (National Laboratory Animal Center, Taiwan) were used for behavioral testing (Figure S1) and an additional recording experiment (Figure S4). During training and recording procedures, these rats were water restricted to their 85-90% weight and were trained in a daily 60-minute session. Water-restricted rats received 15 minutes water access at the end of each training day with free access on weekends.

### Apparatus

Plexiglass operant chambers (11” L×8 ¼” W×13” H), custom-built by Med Associates Inc. (St. Albans, VT), were contained in sound-attenuating cubicles (ENV-018MD) each with an exhaust fan that helped mask external noise. Each chamber’s front panel was equipped with an illuminated nose poke port (ENV-114M) located in the center (horizontal axis) as the fixation port, which was equipped with an infrared (IR) sensor to detect the entry of the animals’ snout into the port. On each side of the center nose-poke port there were two reward ports (CT-ENV-251L-P). Two IR sensors were positioned to detect reward-port entry and sipper-tube licking, respectively.

Sucrose solution (13.3%) was used as reward and delivered through the sipper tubes located in the reward ports. Reward delivery was controlled by solenoid valves (Parker Hannifin Corp #003-0111-900, Hollis, NH) and calibrated to provide 10 µl of solution per drop. Each chamber was equipped with a ceiling-mounted speaker (ENV-224BM) to deliver auditory stimuli, and a stimulus light (ENV-221) positioned above the center fixation port to serve as the new light stimulus. For the additional behavioral testing (Figure S1), one stimulus light each was added above the left and the right reward ports to serve as the sensory cue in the visual discrimination experiment, and water was used as the reward. Behavioral training protocols were controlled by Med-PC software (Version IV or V, Med Associates Inc.), which stored all event timestamps at 1 or 2 ms resolution and also sent out TTL signals to the neurophysiology recording systems.

### Behavioral Training Procedures

Rats were trained in operant chambers that were dimly lit. Rats were first trained in an auditory or visual discrimination task. See TABLE 1 and Figure S1 for details of the stimuli used for each animal. Trials were separated by an unsignaled inter-trial interval (ITI) lasting 4-6 sec. Fixation or licks during the ITI reset the ITI timer. After the ITI, the center fixation port was illuminated, which was turned off only when rats poked the fixation port. Rats were required to maintain fixation in the center nose poke port for a variable amount of foreperiod. Four different foreperiods (0.35, 0.5, 0.65, and 0.8 s) were used, pseudorandomly across trials. After the foreperiod, one of three conditions was randomly presented with equal probabilities: a S^right^ stimulus indicated reward on the right port; a different S^left^ stimulus indicated reward on the left port; or the absence of stimulus (catch trial) indicated no reward. An internal timestamp was recorded in catch trials to mark the onset of the would-be stimulus. Early fixation port exit before the end of the foreperiod led to the re-illumination of the center fixation port. Licking in the correct port within a 3 s window after stimulus onset led to three drops of reward, delivered starting at the 3^rd^ lick. The delivery of reward at the 3^rd^ lick created an expectation for trial outcome at this time point, which was dissociated in time from the initiation of licking (1^st^ lick). We will therefore refer to the time point of the 3^rd^ lick also as the trial outcome event. Trials with licking responses ended after 1 s following the last lick, while trials without licking responses ended after the 3 s response window. The ending of each trial started the ITI timer.

After reaching asymptotic behavioral performance in the auditory or the visual discrimination task, a new learning phase was introduced by replacing either the S^right^ or S^left^ stimuli by a novel sensory stimulus in a different sensory modality, while all other aspects of the task remained the same. In the group that were initially trained with auditory discrimination, the S^right^ sound stimulus was replaced by the central light above the fixation port to indicate reward on the right port (TABLE 1). In the group there were initially trained with visual discrimination, either the S^right^ or S^left^ light stimuli was replaced by a 6 kHz sound (70dB) played from a speaker above the center fixation port (Figure S1).

### Stereotaxic Surgery and Electrode

Surgery was performed under isoflurane anesthesia as previously described (*22*). Multiple skull screws were inserted to anchor the implant, with one screw over the cerebellum serving as the common electrical reference and a separate screw over the opposite cerebellum hemisphere serving as the electrical ground. Craniotomies were opened to target bilateral BF (AP –0.6 mm, ML ±2.25 mm relative to Bregma) (*31*). The electrode contained two bundles of 16 polyimide-insulated tungsten wires (38 µm diameter; California Fine Wire, CA), each bundle ensheathed in a 28-gauge stainless steel cannula and controlled by a precision microdrive. The impedance of individual wire was ~ 0.1 MΩ measured at 1 kHz (niPOD, NeuroNexusTech, MI or Open Ephys Acquisition Board). During surgery, the cannulae were lowered to DV 6.5 mm below cortical surface using a micropositioner (Model 2662, David Kopf Instrument or Robot Stereotaxic, Neurostar GmbH) at a speed of 2-50 µm/s. After reaching target depth, the electrode and screws were covered with dental cement (Hygenic Denture Resin), and electrodes further advanced to 7.5 mm below the cortical surface. Rats received ibuprofen and topical antibiotics after surgery for pain relief and prevention of infection, and were allowed one week to recover with *ad libitum* food and water. Cannulae and electrode tip locations were verified with cresyl violet staining of histological sections at the end of the experiment. All electrodes were found at expected positions between AP [–0.2, –1.2] mm, ML [1.5, 3] mm, relative to Bregma, and DV [7.5, 8.5] mm relative to cortical surface (Figure 3A).

### Data acquisition and spike sorting

Electrical signals were referenced to a common skull screw placed over the cerebellum. Electrical signals were filtered (0.3 Hz–7.5 kHz) and amplified using Cereplex M digital headstages and recorded using a Neural Signal Processor (Blackrock Microsystems, UT). Single unit activity was further filtered (250 Hz–5 kHz) and recorded at 30 kHz. Spike waveforms were sorted offline to identify single units using the KlustaKwik sorting algorithm followed by a custom Python GUI for manual curation. Only single units with clear separation from the noise cluster and with minimal (<0.1%) spike collisions (spikes with less than 1.5 ms interspike interval) were used for further analyses, consistent with previous studies of BF bursting neurons (*21–26*). Additional cross-correlation analysis was used to remove duplicate units recorded simultaneously across multiple electrodes (*21–26*).

### Recording during the new learning phase

After surgery, BF neuronal activity was monitored while rats were re-trained in the auditory discrimination task to asymptotic performance level. During this re-training phase, BF electrode depths were adjusted slightly (by advancing electrodes at 125 µm increment) until a stable population of BF single units can be recorded. At this point, the new learning phase with the light as the new S^right^ stimulus was introduced and rats were trained and recorded daily with BF electrodes remained at the same depth. This approach allowed us to monitor the activity of a large population of BF neurons and follow its temporal evolution across sessions.

## Data analysis

Data were analyzed using custom Matlab (MATLAB The MathWorks Inc., Natick, MA) scripts.

### Define different behavioral response types

Licking responses were defined for stimulus (S^left^ and S^right^) and catch trials if rats licked at least three times in the reward port within the 3 sec window after stimulus onset (or the corresponding timestamp for the would-be stimulus in catch trials). Licking responses to the correct reward port were rewarded with three drops of water, delivered starting at the 3^rd^ lick (referred to as trial outcome event). During the new learning phase, licking responses in the new stimulus and catch trials were predominantly to the reward port associated with the new stimulus.

No-fixation licks corresponded to licking responses to the reward port associated with the new stimulus that were not preceded by poking the center fixation port. Specifically, no-fixation licks were defined based on three criteria: (1) rats made at least three consecutive licks in the reward port; (2) the interval between the last exit from the fixation port and the first lick must be greater than 2 sec; (3) the interval between the last exit from the reward port and the subsequent first lick in the same reward port must be greater than 1 sec. These duration thresholds were determined based on the empirical licking patterns across animals. In the analyses of learning dynamics in the D_1_ session (Figures 3 & 4), no-fixation licks were treated as rightward licking trials, even though such behaviors were self-initiated and not imposed by the task design.

Reaction time (RT) in light trials in a session (Figure 7C) was defined as the median of the interval between the onset of the light stimulus and the exit from the fixation port in light lick trials. Lick duration in catch trials in a session (Figure S3) was defined as the median of the interval between the first and the last lick in catch lick trials.

### Define the D_0_, D_1_ and D_2_ learning landmarks

During the new learning phase, three sessions (D_0_, D_1_, D_2_) were identified in individual animals as landmarks that demarcated distinct stages of new learning (Figure 2C). The D_0_ session was defined as the very first session the new light stimulus was introduced. The D_1_ session was defined as the first session when animals began to respond correctly in the new stimulus trials and obtained reward in the associated reward port in at least three trials. The D_2_ session was defined as the session in which catch licks occurred most frequently. The D_1_ and D_2_ landmarks allowed us to identify similar learning stages across animals despite their individual differences in learning dynamics. The specific timing of the three landmark sessions in each animal are provided in TABLE 2 (also see Figure 2C). One animal (ID#7) with accelerated learning dynamics, in which D_1_ and D_2_ occurred in the same session, was excluded from analyses of D_1_ neural dynamics (Figures 4 & 5) to ensure that neural activities associated with D_2_ did not confound the neural dynamics in the D_1_ session. The BF neuronal activity in this animal was included in the analysis of D_2_ neural dynamics (Figure 6) and showed the strongest phasic response to the light onset among all animals, consistent with its accelerated learning dynamics.

### Identification of the behavioral transition point in the D_1_ session

The behavioral transition points in D_1_ sessions (Figures 3 & 4) were identified based on behavioral response patterns in three trial types combined: light trials, catch trials and no-fixation licks. The behavioral response pattern in each trial was coded as either 1 or 0 based on whether animals licked in the right reward port in that trial. The behavioral transition point was defined as the point with the largest difference in licking responses between the 20 trials before and the 20 trials after that point. In 5/7 animals, the first trial after the behavioral transition was a rewarded light lick trial. In the other two animals, the transition point was adjusted to the closest light lick trial by 2 or 4 trials, respectively.

### Identification of BF bursting neurons

BF bursting neurons were defined as BF single units whose average firing rates during the [0.05, 0.2]s window after stimulus onset increased by more than 2 spikes/s in the S^left^ sound trials compared to the corresponding window in catch trials (Figure S2). This contrast between sound trials and catch trials was necessary because it removed the nonstationary baseline before stimulus onset and allowed us to ask whether BF neurons truly responded to the sound stimulus. In addition, BF bursting neurons should have baseline firing rates (during the [-1, 0]s window relative to the trial start signal) less than 10 spikes/s.

A total of 1453 BF single units were recorded over 45 sessions (N=7 rats), of which 70% (1013/1453) were classified as BF bursting neurons based on their stereotypical phasic response to the S^left^ sound (22.5±7.3 neurons per session, mean±std) (Figure S2, TABLE 2). One session with only one BF bursting neuron was excluded from the analysis of BF population activities. The large number of BF bursting neurons recorded in each session allowed us to treat them as a representative sample of all BF bursting neurons, whose responses to the S^left^ sound were highly stable throughout the learning process (Figure 3B-D). This strategy ensured that we were following functionally the same neuronal ensemble and could track how BF bursting neurons acquired responses to the new light during learning, regardless of whether the identities of these BF neurons were exactly the same in each session.

### Population BF responses to behavioral events

The spike timestamps of all BF bursting neurons in a single session were pooled together to approximate the population activity of all BF bursting neurons. Population peri-stimulus time histograms (PSTHs) were calculated with 10-ms bins, and normalized by the number of BF bursting neurons in a session.

To properly assess whether BF bursting neurons responded to the onset of the new light stimulus (Figures 5-7), it was important to disambiguate such stimulus-onset responses from the increased BF activities after fixation port exit (Figure 5). To achieve this goal, PSTHs to the stimulus onset were calculated based only on spikes that occurred before fixation port exit in individual trials, resulting in different interval lengths (between stimulus onset to fixation port exit) across trials. Accordingly, the calculation of the mean PSTH across all trials in a session was adjusted for the different number of trials at different interval lengths. The mean PSTHs were further truncated at the median interval length of that session to reduce noisy estimates of PSTHs at long interval lengths due to lower number of trials. When PSTHs were averaged across animals, the averaged PSTHs were further truncated at the mean of median interval latencies across animals. This truncation procedure resulted in the uneven lengths of PSTHs across individual animals (Figures 5 & 6) and across sessions (Figure 7). This procedure was also applied to calculating the BF responses before fixation port exit to include only spikes that occurred after stimulus onset (Figures 5 & 6), and for calculating BF responses during licking in catch lick trials (Figure S3).

The time windows used to quantify average BF activity in different epochs were indicated in respective figures, and corresponded to the following: [0.05, 0.2]s after S^left^ sound onset; [0.1, 0.3]s after S^right^ light stimulus onset; [0.1, 0.3]s after the timestamp for the would-be stimulus in catch trials; [0.05, 0.35]s after the 3^rd^ lick for outcome responses; [-0.3, 0]s and [0, 0.3]s relative to the fixation port exit. The epoch for calculating evaluation response is described below.

Evaluation response (Figures 3E, 4B, 7A, 8B, 8C) refers to the increased BF activity after exiting the fixation port and before the trial outcome (3^rd^ lick). The evaluation response reflected animals’ internal evaluation because no additional sensory stimuli were presented during this epoch. Specifically, the evaluation response was calculated in individual trials and defined as the maximum firing rate of any 500ms window during the evaluation epoch, which corresponded to the interval between [fix-out, outcome], with additional adjustments according to trial types. In light lick and catch lick trials, the evaluation epoch was defined as [fix-out, outcome] in each trial. The epoch durations in light lick and catch lick trials within each session were used as the reference point for other trial types as described next. In no-fixation licks, in which the fix-out event was absent, the duration of evaluation epoch was set as the 95^th^ percentile of the evaluation epoch durations in light lick and catch lick trials. In light no lick and catch no lick trials, in which the 3^rd^ lick event was absent, the duration of the evaluation epoch was set as the median of the evaluation epoch durations in light lick and catch lick trials. These adjustments in the definition of evaluation epochs, as well as its calculation of maximum firing rate within the epoch, took into consideration the behavioral variability across trial types, learning stages and individual animals.

To evaluate the dynamic changes of BF activities around the transition point in the D_1_ session (Figures 3E & 4B), single trial evaluation and outcome responses were smoothed using moving median over 10 trials. The smoothed trends were aligned at the transition point and then averaged across all animals. Only trials with smoothed trend data from at least 4 animals were plotted in the group average (Figure 4B).

### Statistics

Statistical comparisons were conducted using the Statistics and Machine Learning Toolbox (version 11.3) in MATLAB (R2018a) (https://www.mathworks.com/). Paired t-test (ttest.m) was used to compare behavioral and neural activity differences between two groups (Figures 3D, 6A, 7A). Repeated measures analysis of variance (ranova.m) was used for comparisons involving more than 2 groups, by specifying the appropriate within-subject models (Figures 4A3, 4B3, 5B). Comparisons of PSTHs between two groups (Figures 5A, 6B, 6C) was conducted for each 100 ms sliding window (10 ms step) using paired t-test. Significance level was set at p<0.01 for three consecutive bins. Pearson correlation (corrcoef.m) was used to determine the relationship between neuronal activities and/or behavior (Figures 7, 8, S3B).

### Receiver operating characteristic (ROC) and area under curve (AUC) analysis

To determine whether the activity of BF bursting neurons differentiated between trial types within each D_1_ session (Figure 5C), we compared BF activity for each 100ms sliding window (10ms step) using the AUC measure of ROC analysis (auc.m by Alois Schloegl). At each sliding window, BF population activity was calculated for each light and catch trial, and distributions of BF activities were compared between light vs catch trials or between lick vs no lick trials. Significance level was set at p<0.001 using 10,000 trial-shuffled random permutations.

To determine whether BF activity differentiated between lick and no lick trials within the same trial type (light or catch trials) (Figure S3A), we compared BF activity in the [0, 500] ms window after exiting the fixation port. For each session and each trial type, lick and no lick trials must each constitute at least 10% of that trial type to be included in the analysis. Catch trials from all sessions were included in this analysis. Only light trials before the D_2_ session (pre-D_2_) were included in this analysis because BF responses to the onset of the light stimulus had not developed in those sessions. Significance level was set at p<0.05 using 1,000 trial-shuffled random permutations.

## Acknowledgements

We thank P.R. Rapp, A. Scaglione, S.W. Wu and K.L. Hsieh for critical discussions of the manuscript; B.M. Brock for technical support. This research was funded by the Intramural Research Program of the National Institute on Aging (NIH, USA) and by NARSAD Young Investigator Award to S.L.. Additional supports came from the National Science and Technology Council (NSTC, Taiwan) grants (107-2320-B-010-028-MY3; 108-2638-B-010-002-MY2; 110-2628-B-A49A-505), and from the Brain Research Center, National Yang Ming Chiao Tung University from The Featured Areas Research Center Program within the framework of the Higher Education Sprout Project by the Ministry of Education (MOE) in Taiwan.

## Author Contributions

H.E.M. and S.L. designed the study. H.E.M., K.V., Y.J. and H.C. performed experiments and collected data. H.E.M. and S.L. analyzed data. H.E.M. and S.L. wrote the manuscript with inputs from K.V., Y.J., and H.C.

## Supplementary Figures

**Figure S1.**
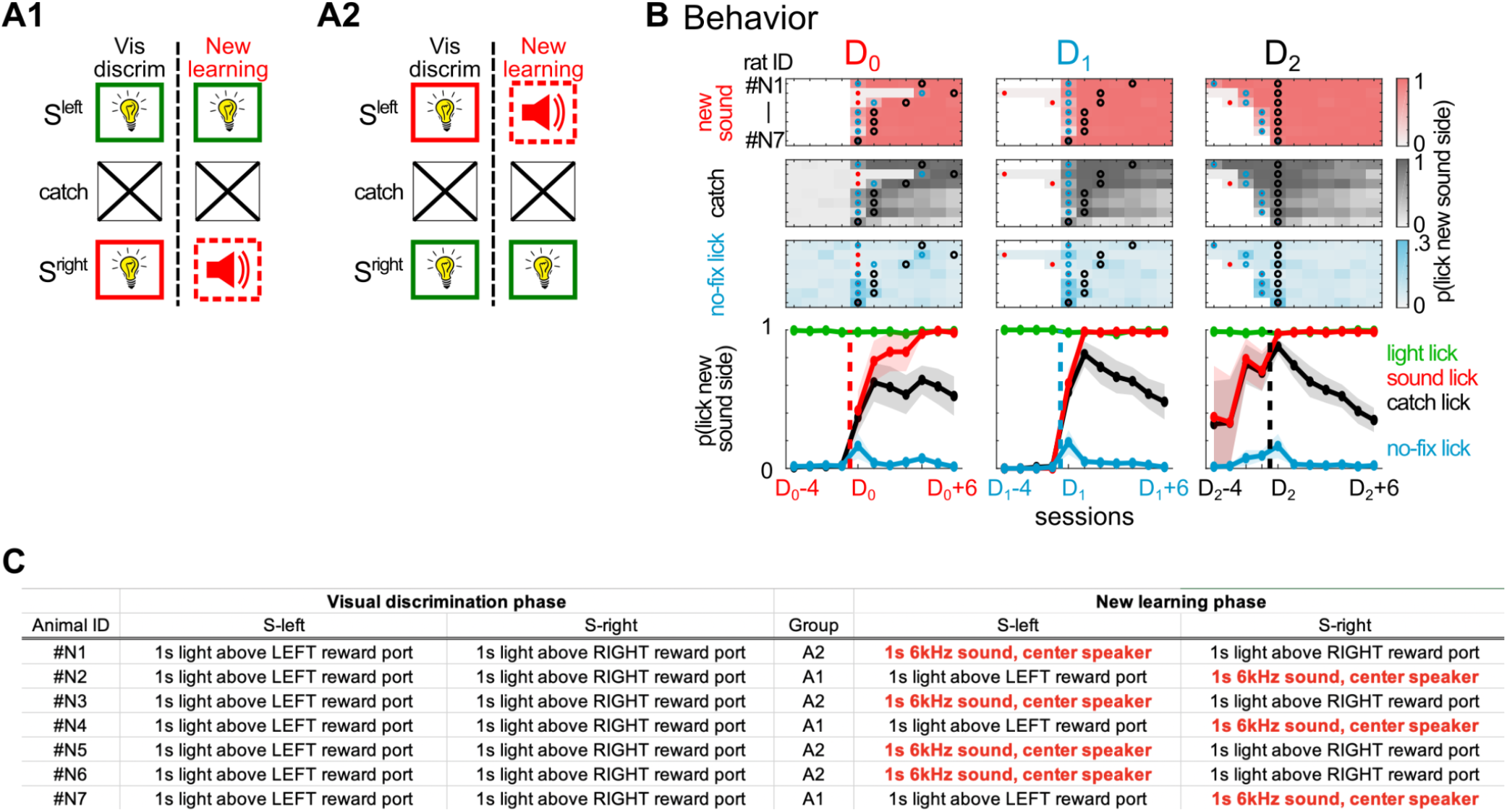
Consistent behavioral response patterns during new learning under variations of stimulus modalities and response directions. **A**, A separate cohort of 7 rats were divided into two groups. Both groups initially learned a visual discrimination task, with lights above the left or right reward port serving as the sensory cue for reward in the respective port. At the new learning phase, one of the two lights was replaced by a 6 kHz (70dB) sound, delivered from a speaker placed above the center fixation port. In the A1 group, the S^right^ light was replaced (N=3); while in the A2 group, the S^left^ light was replaced (N=4). All other elements of the task remained the same. **B**, The proportion of the three types of reward-seeking behaviors toward the reward port associated with the new sound stimulus side during new learning (N=7 rats). Conventions as in Figure 2C. Similar to the results in Figure 2C, no-fixation licks were most consistently associated with the D_1_ session, which occurred earlier than the peak of catch licks (D_2_ session). Further, the proportion of sound licks and catch licks were highly similar prior to the D_2_ session, and began to diverge at the D_2_ session. Thus, the pattern of sequential refinements can be observed regardless of the sensory modality of the stimulus or the laterality of the new learning side. **C**, Stimulus parameters of the S^left^ and S^right^ stimuli for each animal in this cohort.

**Figure S2.**
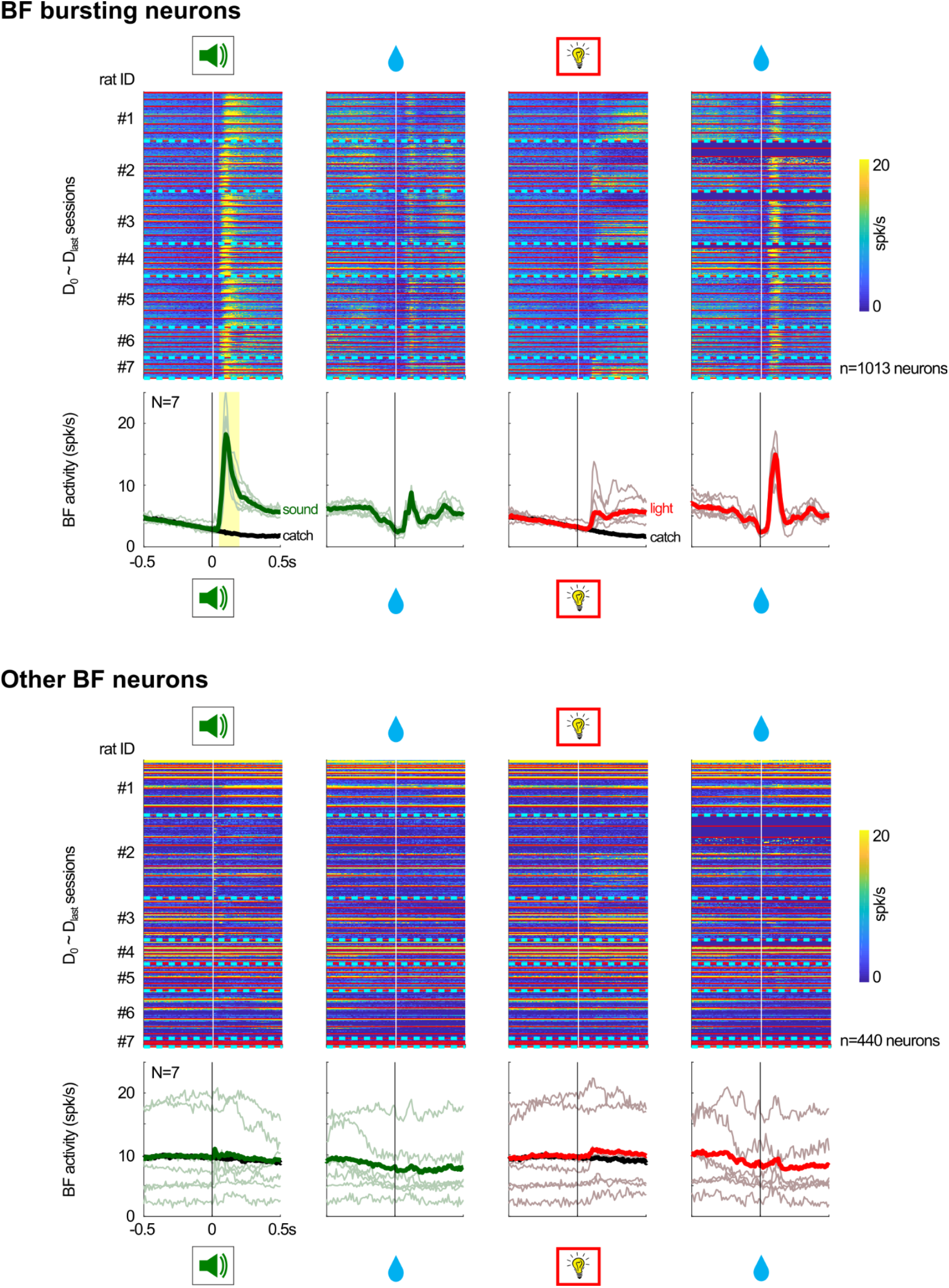
Identification of BF bursting neurons and their responses to behavioral events. Responses of individual BF bursting neurons (n=1013) and other BF neurons (n=440) to behavioral events (stimulus onset and reward) in the S^left^ sound trials (two left columns) and in the S^right^ light trials (two right columns). Recordings from different sessions (N=45 sessions; separated by thin red lines) and different animals (N=7 rats; separated by cyan dotted lines) were separated by horizontal lines. Lower panels showed the average response (thick lines) pooled across individual animals (thin lines). BF activities in catch trials (thick black lines) were plotted for comparison. Responses to the stimulus onset event were calculated based on all trials in that session, regardless of subsequent behavioral responses (licking or not). On the other hand, responses to the reward were calculated based only on correct licking trials. Conventions as in Figure 3B-C. BF bursting neurons were defined as BF single units whose average firing rates during the [0.05, 0.2]s window after stimulus onset (yellow shaded interval) increased by more than 2 spikes/s in the S^left^ sound trials compared to the corresponding window in catch trials. This contrast between sound trials and catch trials was necessary because many BF neurons changed their activity during the foreperiod while waiting for stimulus onset. In addition, BF bursting neurons should have baseline firing rates less than 10 spikes/s. The activities of BF bursting neurons in S^left^ sound trials were highly similar across sessions and across animals (Figure 3C-D). Note that the calculation of these PSTHs to the stimulus onset event (as well as those in Figure 3B-D) did not exclude spikes that occurred after fixation port exit (as in Figures 5-7). The truncation procedure used in Figures 5-7 was needed to disambiguate BF responses to light onset from the increased BF activities after fixation port exit (i.e. evaluation responses).

**Figure S3.**
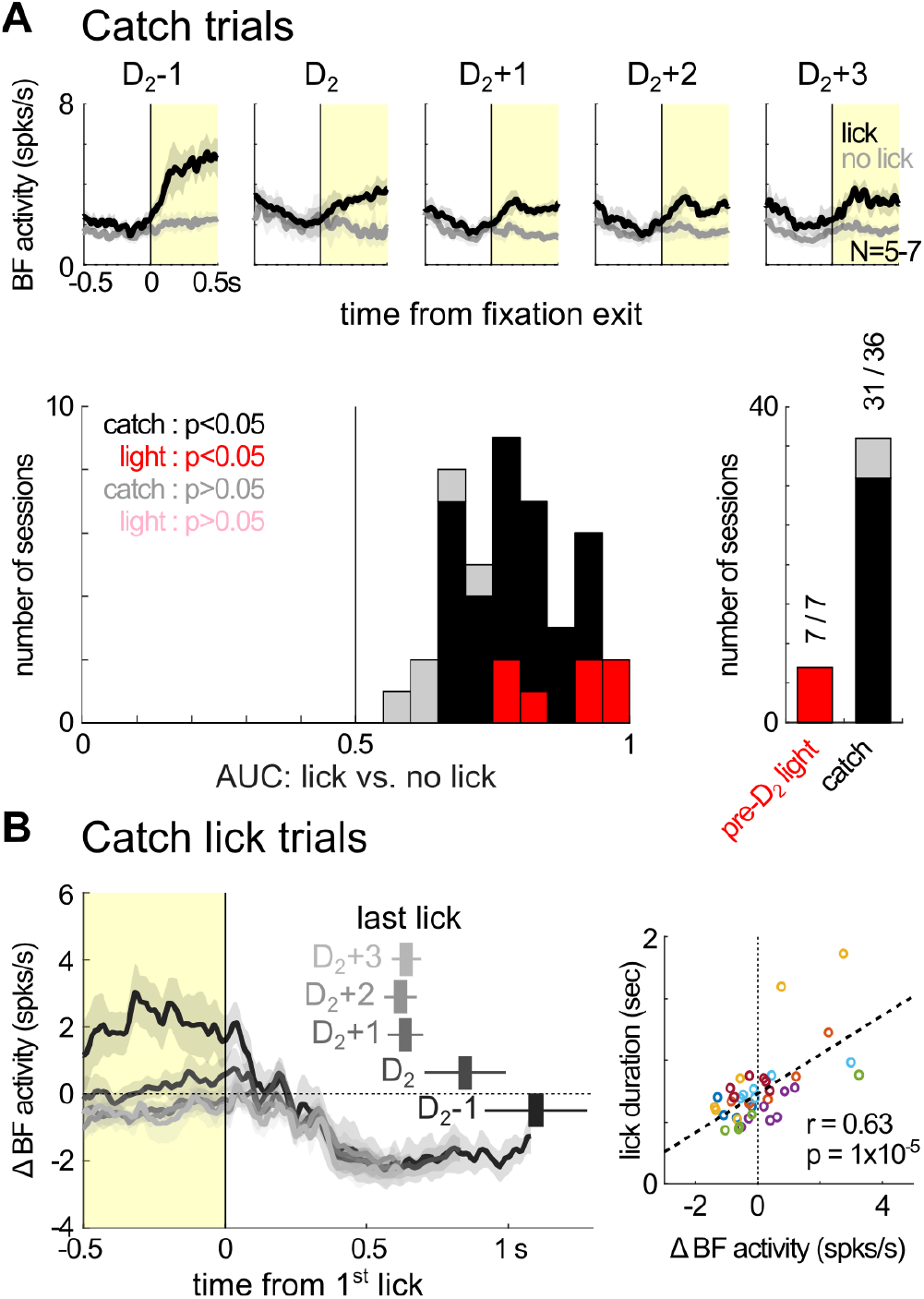
Increased BF activity predicted reward-seeking behaviors in the absence of the light stimulus. **A**, Average responses of BF bursting neurons (mean ± s.e.m.) in catch trials aligned at fixation port exit, plotted separately for lick and no lick trials (top). Distributions of AUC values from comparing BF activities in the [0, 0.5]s window after fixation port exit (yellow shaded interval) between lick and no lick trials within the same trial type (catch or light trials) (bottom). Within the same trial type, increased BF activities reliably predicted reward-seeking behavior toward the right reward port. Only light trials from pre-D_2_ sessions were included because BF responses to the light stimulus had not developed. **B**, Average responses of BF bursting neurons (mean ± s.e.m.) in catch lick trials aligned at the first lick (left). BF activities, relative to their respective baseline firing rates, were truncated at the median lick duration of the respective sessions. The timing of the last lick (mean ± s.e.m.) of the corresponding sessions were shown above each trace. BF activities prior to the start of licking in catch lick trials (yellow shaded interval, left panel) were positively correlated with the median lick duration in individual sessions (right). Each circle indicates one session and different colors correspond to different animals.

**Figure S4.**
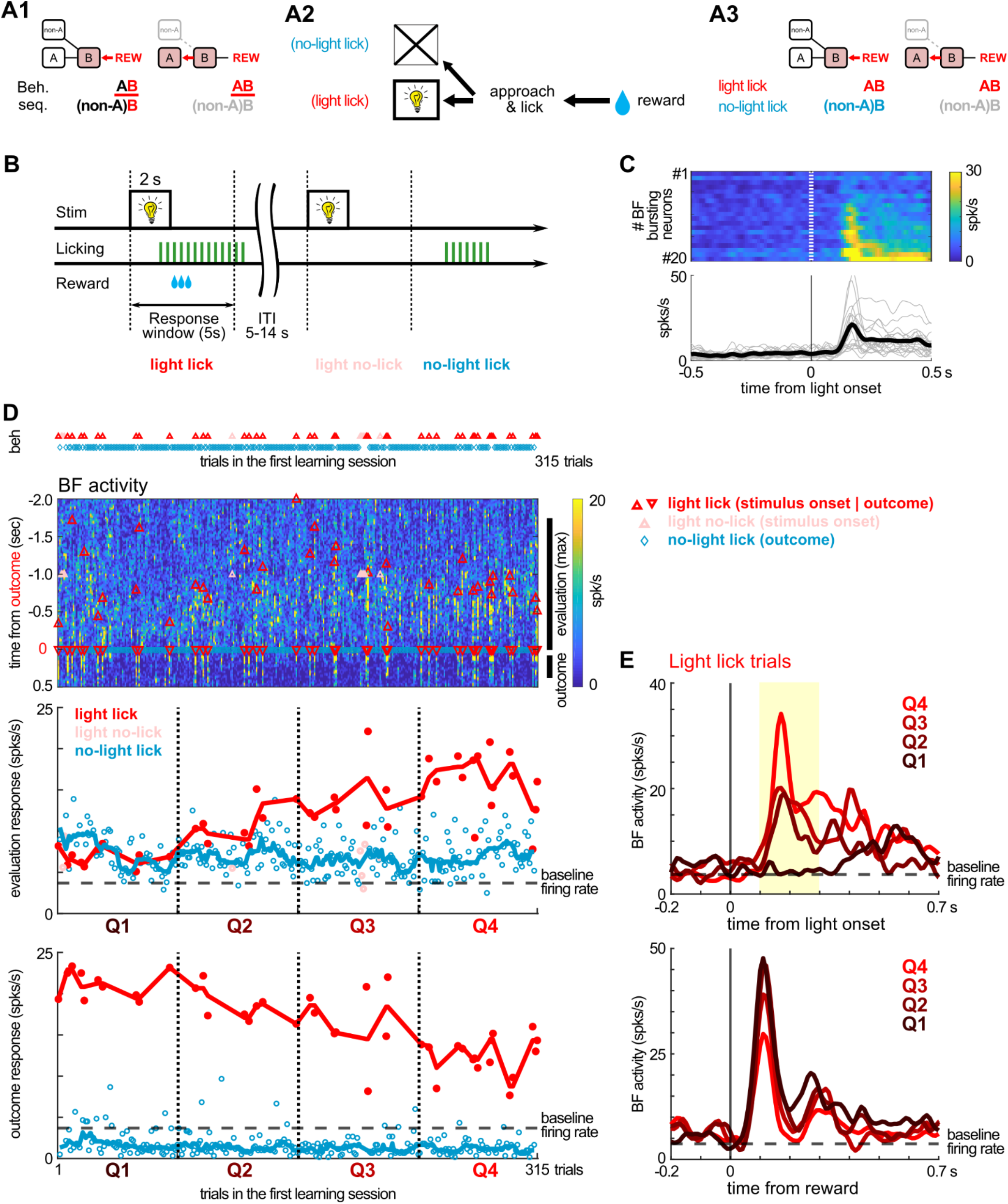
Stepwise learning of a two-element behavioral sequence. **A**, The conceptual model of how a two-element sequence is learned using the stepwise strategy. **A1**, A model depicting the two steps in learning a two-element (A-B) behavioral sequence. Conventions as in Figure 1B. **A2**, In this experiment, the A element corresponded to the light stimulus, and B the approach and licking response. Only the A-B sequence was rewarded. **A3**, The model predicts that, during the first step of learning, two reward-seeking behavioral sequences - light licks and no-light licks - will be observed, and both should be accompanied by similar levels of BF activities. **B**, New learning task. House light served as the reward-predicting stimulus. Licking at the reward port within a 5-sec response window led to water delivery starting at the third lick. Inter-trial interval (ITI) was randomly chosen between 5-14 sec. Licking in the absence of the light stimulus was not rewarded and reset the ITI counter. Three types of behaviors - light licks, light no-licks, no-light licks - were depicted in this schematic. No-light lick was defined as a licking cluster of at least 3 licks in length, with its first lick occuring outside the 5 sec response window of light onset, and at least 3 sec after the end of the last lick cluster. **C**, The average responses of BF bursting neurons to light onset during the first session of new learning in one example rat. 20/29 BF neurons recorded in this session were classified as BF bursting neurons based on two criteria: (1) BF activities in the [0.1, 0.3]s window after light onset increased by 2 spikes/s over baseline firing rates; (2) Baseline firing rates were within [0.1, 10] spikes/s. **D**, Behavioral and BF neuronal responses during the first learning session, plotted against trial sequence (x-axis) in this session (315 trials). **Top panel**, behavioral responses across trials. **Second panel**, population activities of BF bursting neurons across the same trial sequence (x-axis). Y-axis indicates time in each trial, with time zero aligned at the trial outcome (defined as the timing of the third lick). Light no-lick trials were aligned instead at the time of stimulus onset (pink triangles) such that the median timing of light onset in light lick and light no-lick trials were equivalent. The black lines to the right of the panel indicate the time windows for calculating evaluation and outcome responses. **Third panel**, BF evaluation responses, plotted separately for the three trial types. Evaluation response was calculated as the maximum firing rate of any 500ms window during [-1.75, 0]s before trial outcome. Circles indicate BF activities in single trials and lines indicate their respective trends (10-trial moving medians). Note that during the first quartile of trials (Q1), evaluation responses were similar between light licks and no-light licks, which became distinct in later quartiles. **Fourth panel**, BF outcome responses across trials, plotted separately for the two trial types with licking behaviors. Outcome response was calculated as the mean BF activity during [0.05, 0.35]s after the 3^rd^ lick. Outcome responses to reward delivery became weaker over trials. **E**, Average BF responses to light onset and reward delivery in the four trial quartiles. Notice that during Q1, there was no phasic BF response to light onset. This suggests that the increased BF evaluation responses in Q1 (third panel in D) did not result from BF responses to the light onset, and instead reflected an internal evaluation signal that was similarly present in both light licks and no-light licks. This observation supports the stepwise learning strategy (panel A), with Q1 trials corresponding to the first step of learning, during which animals engaged in two types of reward-seeking behaviors - light licks and no-light licks. Light licks in Q1 trials were not driven by the light stimulus. Learning about the light stimulus occurred in later trials when BF responses to light onset emerged, which was also when BF evaluation responses began to diverge between light licks and no-light licks.

## References

1. I. P. Pavlov, Lectures on conditioned reflexes: Twenty-five years of objective study of the higher nervous activity (behaviour) of animals (Liverwright Publishing Corporation, New York, 1928), vol. 414.

2. R. A. Rescorla, A. R. Wagner, in Classical conditioning: current research and theory (1972), vol. 2.

3. E. L. Thorndike, The Psychological Review: Monograph Supplements, in press.

4. P. Dayan, B. W. Balleine, Reward, motivation, and reinforcement learning. Neuron. 36, 285–298 (2002).

5. B. F. Skinner, The behavior of organisms: an experimental analysis (Appleton-Century, Oxford, England, 1938), vol. 457.

6. J. P. O’Doherty, A. Hampton, H. Kim, in Annals of the New York Academy of Sciences (2007), vol. 1104, pp. 35–53.

7. B. B. Doll, D. A. Simon, N. D. Daw, The ubiquity of model-based reinforcement learning. Curr. Opin. Neurobiol. 22, 1075–1081 (2012).

8. E. C. Tolman, Cognitive maps in rats and men. Psychol. Rev. 55, 189–208 (1948).

9. R. C. Wilson, Y. K. Takahashi, G. Schoenbaum, Y. Niv, Orbitofrontal cortex as a cognitive map of task space. Neuron. 81, 267–279 (2014).

10. T. E. J. Behrens, T. H. Muller, J. C. R. Whittington, S. Mark, A. B. Baram, K. L. Stachenfeld, Z. Kurth-Nelson, What Is a Cognitive Map? Organizing Knowledge for Flexible Behavior. Neuron. 100, 490–509 (2018).

11. W. Schultz, P. Dayan, P. R. Montague, A neural substrate of prediction and reward. Science. 275, 1593–1599 (1997).

12. N. Eshel, M. Bukwich, V. Rao, V. Hemmelder, J. Tian, N. Uchida, Arithmetic and local circuitry underlying dopamine prediction errors. Nature. 525, 243–246 (2015).

13. M. Pessiglione, B. Seymour, G. Flandin, R. J. Dolan, C. D. Frith, Dopamine-dependent prediction errors underpin reward-seeking behaviour in humans. Nature. 442, 1042–1045 (2006).

14. R. S. Sutton, A. G. Barto, Reinforcement Learning: An Introduction. IEEE Trans. Neural Netw. 9, 1054–1054 (1998).

15. D. Hassabis, D. Kumaran, C. Summerfield, M. Botvinick, Neuroscience-Inspired Artificial Intelligence. Neuron. 95, 245–258 (2017).

16. B. A. Richards, T. P. Lillicrap, P. Beaudoin, Y. Bengio, R. Bogacz, A. Christensen, C. Clopath, R. P. Costa, A. de Berker, S. Ganguli, C. J. Gillon, D. Hafner, A. Kepecs, N. Kriegeskorte, P. Latham, G. W. Lindsay, K. D. Miller, R. Naud, C. C. Pack, P. Poirazi, P. Roelfsema, J. Sacramento, A. Saxe, B. Scellier, A. C. Schapiro, W. Senn, G. Wayne, D. Yamins, F. Zenke, J. Zylberberg, D. Therien, K. P. Kording, A deep learning framework for neuroscience. Nat. Neurosci. 22, 1761–1770 (2019).

17. B. F. Skinner, The Reinforcing Effect of a Differentiating Stimulus. J. Gen. Psychol. 14, 263–278 (1936).

18. M. Enquist, J. Lind, S. Ghirlanda, The power of associative learning and the ontogeny of optimal behaviour. R Soc Open Sci. 3, 160734 (2016).

19. S. Ghirlanda, J. Lind, M. Enquist, A-learning: A new formulation of associative learning theory. Psychon. Bull. Rev. 27, 1166–1194 (2020).

20. P. McGreevy, R. Boakes, Carrots and Sticks: Principles of Animal Training (Darlington Press, 2011).

21. S.-C. Lin, M. A. L. Nicolelis, Neuronal ensemble bursting in the basal forebrain encodes salience irrespective of valence. Neuron. 59, 138–149 (2008).

22. I. Avila, S.-C. Lin, Motivational salience signal in the basal forebrain is coupled with faster and more precise decision speed. PLoS Biol. 12, e1001811 (2014).

23. J. D. Mayse, G. M. Nelson, I. Avila, M. Gallagher, S.-C. Lin, Basal forebrain neuronal inhibition enables rapid behavioral stopping. Nat. Neurosci. 18, 1501–1508 (2015).

24. S. M. Raver, S.-C. Lin, Basal forebrain motivational salience signal enhances cortical processing and decision speed. Front. Behav. Neurosci. 9, 277 (2015).

25. I. Avila, S.-C. Lin, Distinct neuronal populations in the basal forebrain encode motivational salience and movement. Front. Behav. Neurosci. 8, 421 (2014).

26. D. P. Nguyen, S.-C. Lin, A frontal cortex event-related potential driven by the basal forebrain. Elife. 3, e02148 (2014).

27. D. J. Ottenheimer, B. A. Bari, E. Sutlief, K. M. Fraser, T. H. Kim, J. M. Richard, J. Y. Cohen, P. H. Janak, A quantitative reward prediction error signal in the ventral pallidum. Nat. Neurosci. 23, 1267–1276 (2020).

28. D. J. Ottenheimer, K. Wang, X. Tong, K. M. Fraser, J. M. Richard, P. H. Janak, Reward activity in ventral pallidum tracks satiety-sensitive preference and drives choice behavior. Sci Adv. 6 (2020), doi:10.1126/sciadv.abc9321.

29. W. Schultz, Dopamine reward prediction-error signalling: a two-component response. Nat. Rev. Neurosci. 17, 183–195 (2016).

30. S.-C. Lin, R. E. Brown, M. G. Hussain Shuler, C. C. H. Petersen, A. Kepecs, Optogenetic Dissection of the Basal Forebrain Neuromodulatory Control of Cortical Activation, Plasticity, and Cognition. J. Neurosci. 35, 13896–13903 (2015).

31. G. Paxinos, C. Watson, The rat brain in stereotaxic coordinates (Academic Press, London, ed. 6, 2007).

